# Recycling limits the lifetime of actin turnover

**DOI:** 10.1101/2022.09.30.510257

**Authors:** Alexandra Colin, Tommi Kotila, Christophe Guérin, Magali Orhant-Prioux, Benoit Vianay, Alex Mogilner, Pekka Lappalainen, Manuel Théry, Laurent Blanchoin

## Abstract

Intracellular organization is largely mediated by the actin turnover. Cellular actin networks consume matter and energy to sustain their dynamics, while maintaining their appearance. This behavior, called ‘dynamic steady state’, enables cells to sense and adapt to their environment. However, how structural stability can be maintained during the constant turnover of a limited actin monomer pool is poorly understood. To answer this question, we developed an experimental system using actin bead motility in a compartment with a limited amount of monomer. We used the speed and the size of the actin comet tails to evaluate the system’s monomer consumption and its lifetime. We established the relative contribution of actin assembly, disassembly and recycling for a bead movement over tens of hours. Recycling mediated by cyclase-associated proteins is the key step in allowing the reuse of monomers for multiple assembly cycles. Energy supply and protein aging are also factors that limit the lifetime of actin turnover. This work reveals the balancing mechanism for long-term network assembly with a limited amount of building blocks.

## Introduction

Living organisms depend on maintaining a ceaseless renewal of their internal organization in the face of constant environmental changes. This energy-dependent process can be observed at several hierarchical levels, where the stability of a biological component is supported by the turnover of its building blocks (Rafelski & Marshall, 2008; Chan & Marshall, 2012; Goehring & Hyman, 2012). In eukaryotes, the actin cytoskeleton plays many key roles in maintaining dynamic intracellular organization (Chhabra & Higgs, 2007; Lappalainen *et al*, 2022). For example, actin dynamics in the cortex (Fritzsche *et al*, 2013, 2016), lamellipodium (Pollard & Borisy, 2003) or stress fibers (Hotulainen & Lappalainen, 2006; Tojkander *et al*, 2015; Nishimura *et al*, 2021) underpin cell migration (Lai *et al*, 2008; Burnette *et al*, 2011; Rottner & Stradal, 2011; Blanchoin *et al*, 2014), and organelle dynamics (Chakrabarti *et al*, 2021). During these processes, actin filament networks constantly assemble and disassemble, often while maintaining an apparent stable structure necessary for force generation (Blanchoin *et al*, 2014; Lappalainen *et al*, 2022), suggesting that the two processes are precisely balanced. Such behaviors are called “dynamic steady-states”, conferring upon cytoskeletal networks a high degree of plasticity that allows cells to adapt and optimize their architecture in response to external changes (Mueller *et al*, 2017; Banerjee *et al*, 2019; Rafelski & Theriot, 2004; Vargas *et al*, 2016; Lomakin *et al*, 2015). A three-step cycle of assembly, disassembly and recycling (Figure 1A) regulates actin dynamics in cells, which is sustained by constant energy consumption. In step 1, actin monomers loaded with ATP assemble into actin filaments. Prior to disassembly, actin subunits in an actin filament hydrolyze their bound ATP and dissociate the inorganic phosphate product, with the subunit remaining ADP-bound (Pollard *et al*, 2000). In step 2, the ADP-bound subunit dissociates from the filament. In step 3, the depolymerized subunit re-loads with ATP, recycling the subunit for another round of assembly (Plastino & Blanchoin, 2019). One question that remains unanswered is how a dynamic steady state can be set-up and maintained with the limited amount of components available inside the cell cytoplasm.

**Figure 1.**
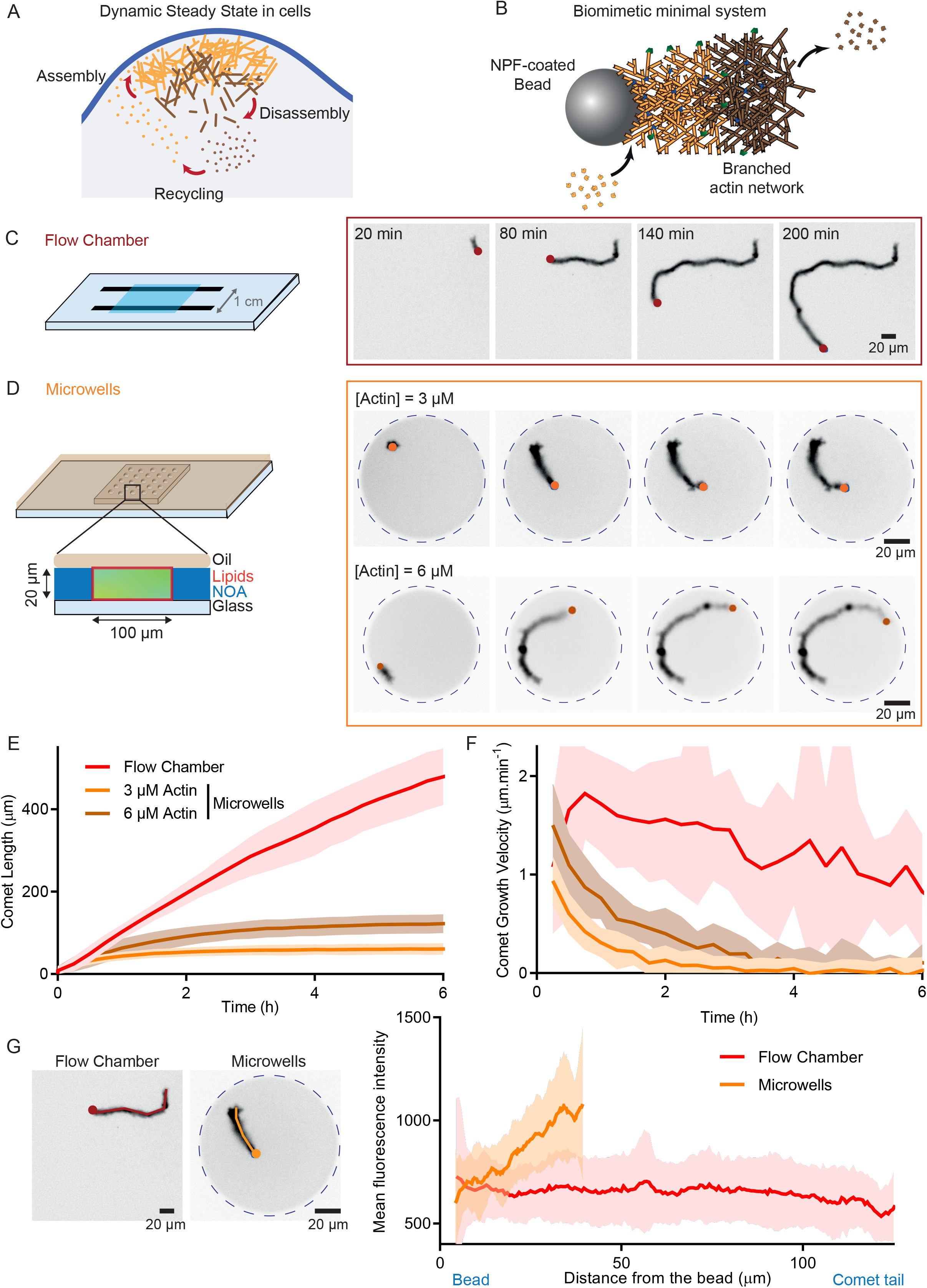
Effect of a limited pool of monomers on actin assembly. **A**. Cartoon of the three-step cycle for actin dynamic steady state in cells. **B**. Schematic of a bead coated with a Nucleation Promoting Factor (NPF) of the Arp2/3 complex able to generate a branched actin comet. Actin assembly takes place on the bead. Disassembly occurs within the tail of the comet. **C**. Left: Scheme of the flow chamber (unlimited amount of proteins) used in this study. Right : Snapshots of the growth of one actin comet from a bead (red dot) in a flow chamber. **D**. Left: Schematic of the microwells used in this study. Right : Snapshots of the growth of one actin comet from a bead (orange) in a microwell for two different initial actin monomers concentration (3 and 6 μM). **E**. Length of comets grown as a function of time in bulk and in microwells. **F**. Growth velocity of comets grown in flow chamber and in microwells as a function of time. Flow chamber: N = 2, n = 33 comets tails. Microwells 3 μM actin: N = 3, n = 38 comets tails. Microwells 6 μM actin: N = 1, n = 17 comet tails. **G**. Mean intensity profiles along comets grown in flow chambers and in microwells (Flow chamber: N = 1, n = 15 comets tails. Microwells 3 μM actin: N = 1, n = 21 comets tails). **Biochemical conditions:** 4.5 μm beads coated with 400 nM of Strep-SNAP-WA. Reaction mix: [Actin] = 3 or 6 μM. [Profilin] = 6 or 12 μM. [Arp2/3 complex] = 90 nM. [Capping Protein] = 15 nM.

*In vitro* experiments, based on a minimal set of purified actin-binding proteins, have dramatically enhanced our understanding of actin network dynamics, with the molecular insights obtained from these approaches complementing *in vivo* experiments. Such experiments have revealed the basics of actin network assembly by the Arp2/3 complex (Mullins *et al*, 1998), actin-based motility and force generation (Loisel *et al*, 1999; Bernheim-Groswasser *et al*, 2002), selectivity in actin network contraction (Reymann *et al*, 2012) or disassembly (Gressin *et al*, 2015), actin network competition (Suarez *et al*, 2015; Antkowiak *et al*, 2019), as well as cytoskeletal network crosstalk (Lopez *et al*, 2014; Henty-Ridilla *et al*, 2016; Colin *et al*, 2018; Alkemade *et al*, 2022). However, these systems rarely reach the dynamic steady-state that occurs in living cells, with networks either growing then stalling or disassembling. Some systems have improved upon this situation by reaching a balance between assembly and disassembly. For example, dynamic actomyosin networks were reconstituted *in vitro* close to a lipid bilayer (Sonal *et al*, 2019) or in *Xenopus* egg extracts encapsulated in oil droplets (Pinot *et al*, 2012; Tan *et al*, 2018; Malik-Garbi *et al*, 2019). Those studies demonstrated that robust and tunable actin flows are regulated by two main components: the actin turnover rate and the network geometry. In parallel, reconstitution of dynamic networks with a balance between assembly and disassembly mediated by actin polymerization and disassembly by ADF/cofilin (Michelot *et al*, 2007; Akin & Mullins, 2008; Reymann *et al*, 2011; Manhart *et al*, 2019; Pollard *et al*, 2020) has allowed to better understand the role of the different molecular actors in the establishment of a dynamic steady state. However, those experiments were performed either with cell extracts where precise protein content is unknown or with purified proteins in unlimited volumes (therefore masking some crucial steps existing when a limited amount of component is available).

Indeed, one major challenge in investigating actin turnover and its impact on dynamic steady-state in reconstituted systems is related to the number of available active molecular constituents over time. *In vitro* experiments use biologically-relevant concentrations of the components but they are typically performed in relatively large volumes. Therefore, in a large environment, the reservoir of components necessary for actin turnover will be unlimited. As a result, depletion of some components, a critical effect in small volumes like that of a cell, cannot be controlled or even accomplished, shifting thus the chemical kinetics to a non-biological regime in which the lifetime of a self-sustaining system cannot be studied. A growing number of studies have used cell-sized compartment such as water in oil droplets, GUVs (Giant unilamellar vesicles) or microchambers in order to mimic the cell volume and to evaluate the effect of biochemical or mechanical parameters on the cytoskeleton self-organization (Jia & Schwille, 2019; Soares e Silva *et al*, 2011; Alvarado *et al*, 2014; Hsu *et al*, 2022; Miyazaki *et al*, 2015; Bashirzadeh *et al*, 2021). However, how cytoskeletal dynamics can be set-up and maintained in these protein-pool-limited environments has been little studied. More precisely the balance between assembly, disassembly and recycling fluxes, coupled with energy supply to maintain dynamic actin structures remains poorly understood.

Here we ask how actin network assembly can be achieved over time in a cell-sized compartment where subunit recycling is required to maintain its limited pool. To address this issue, we developed a novel experimental system that combines actin bead motility and microfabricated microwells. In our cell-sized compartment, we first investigated the minimal biochemical conditions for establishing a sustained actin turnover over time. We quantified how assembly, disassembly and recycling individually or collectively support the maintenance of a dynamic and stable actin structure over long periods of time. Fast recycling by cyclase-associated protein plays a critical role in maintaining the pool of polymerizable actin monomers. This step depends on ATP concentration. Furthermore, we find that even in excess of ATP, the system loses its effectiveness overtime because aging of actin monomers limits the lifetime of actin turnover.

## Results

### Microwells are closed environments where actin assembly is limited over time

To study actin network dynamic, we used the well-described bead motility assay, in which 4.5 μm diameter polystyrene beads are coated with an Arp2/3 complex-activating Nucleation Promoting Factor (NPF, Snap-Streptavidin-WA-His, in this study) (Bernheim-Groswasser *et al*, 2002; Cameron *et al*, 1999, 2001; Akin & Mullins, 2008; Reymann *et al*, 2011). When these beads are introduced to a medium containing the Arp2/3 complex, actin, profilin and capping protein (assembly conditions), an actin comet assembles and beads exhibit directional motility that can be sustained for hours (Figure 1B, 1C, Movie 1).

To evaluate the effect of a limited amount of components on the lifetime of bead-induced actin assembly, we used microwells (similar to the ones described in (Yamamoto *et al*, 2022)). They have a diameter of 100 μm and a height of 20 μm giving an approximate volume of 140 pL, five orders of magnitude smaller than the 20 μl volume of a classical flow chamber used in *in vitro* assays (Figure 1D, Supplemental Figure 1A). The microwells were closed with oil and their internal surface passivated with lipids (Supplemental Figure 1A). The hermeticity of the microwells has been confirmed using FRAP experiment (Supplemental Figure 1B, 1C). We validated that actin assembly is preserved inside microwells by measuring the association rate constant at the actin barbed end (k_+_) and comparing it to actin assembly in bulk (Supplemental Figure 1D). Finally, we also confirmed that the Arp2/3 complex machinery was functional (Supplemental Figure 1E). In all experiments, the time 0 of the reaction was when actin monomers were added to the mix before introducing it into the microwells and sealing them with oil.

**Figure 2.**
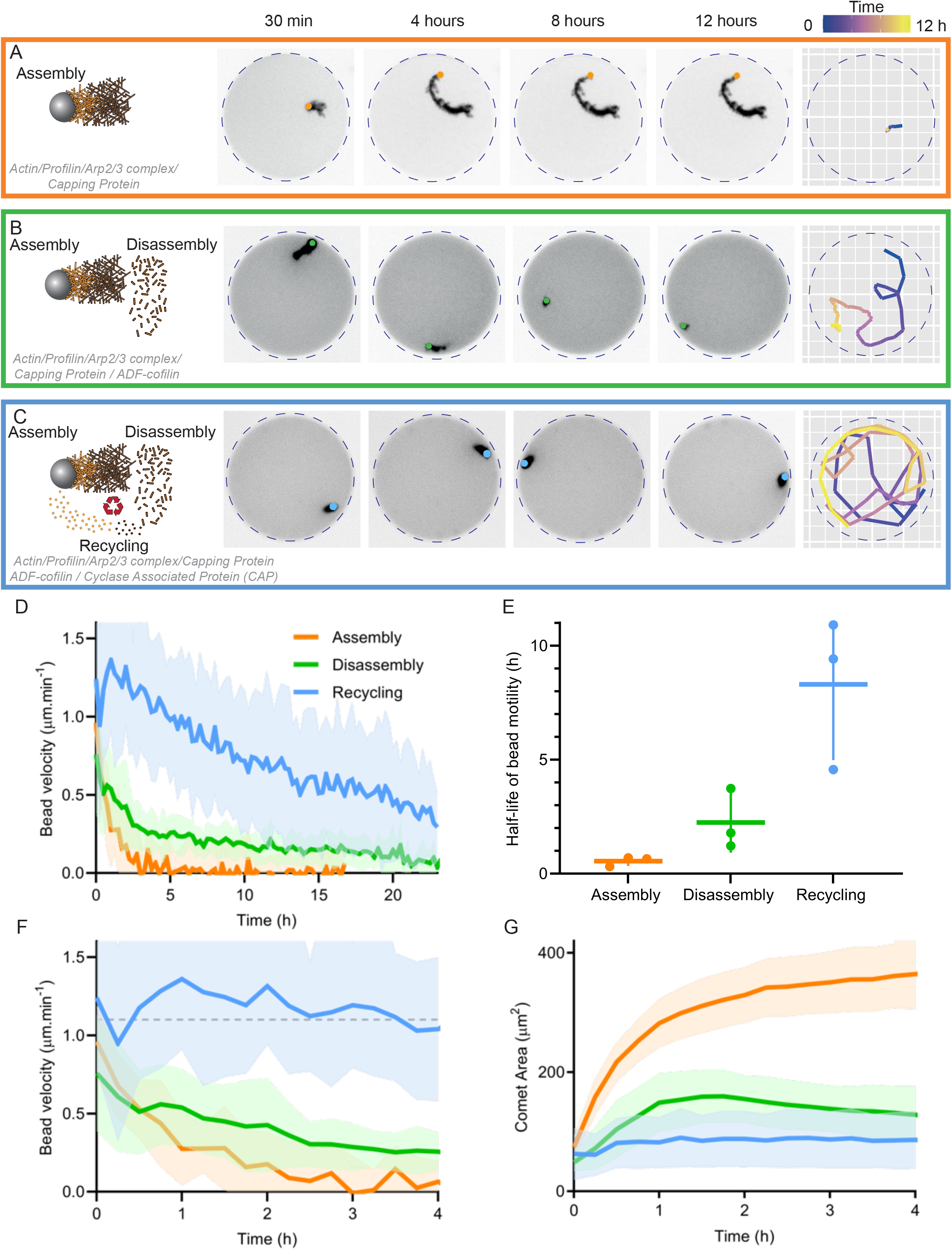
Contribution of assembly, disassembly and recycling on actin turnover in cell-sized compartment. **A**. Left: Snapshots of actin comet tails assembled in Assembly conditions: 4.5 μm polystyrene beads coated with 400 nM Strep-SNAP-WA; 3 μM actin, 6 μM profilin, 90 nM Arp2/3 complex, 15 nM capping protein. Right: Tracking of the comet shown in the snapshots. Time is encoded in color. **B**. Left: Snapshots of actin comet tails assembled in Disassembly conditions: same as assembly conditions with 200 nM ADF/cofilin added. Right: Tracking of the comet shown in the snapshots. Time is encoded in color. **C**. Left: Snapshots of actin comet tails assembled in Recycling conditions: Disassembly conditions with 400 nM Cyclase Associated Protein (CAP) added. Right: Tracking of the comet shown in the snapshots. Time is encoded in color. **D**. Bead velocity as a function of time for one dataset per condition. Mean and standard deviation are represented. **E**. Estimation of half-life of bead motility for each independent dataset in the different dynamic conditions. **F**. Same panel as panel C on a 4 hour time scale. **G**. Comet Area as a function of time for one dataset per condition. Mean and standard deviation are represented. Assembly: n = 20 comet tails. Disassembly: n = 13 comet tails. Recycling: n = 16 comet tails.

We first compared single bead movement in a flow chamber (infinite volume) and in the microwells (cell-sized volume) (Figure 1C-D, Movie 1). In the flow chamber, the length of the comet increases linearly for ∼6 hours while it reached a plateau after ∼3 hours in microwells (Figure 1E). In consequence, the growth velocity of comet tails was constant over time in the flow chamber but decreases rapidly in microwells (Figure 1F). This suggests that in microwells the pool of actin monomers is limited and that after 3 hours most of the monomers were consumed. Interestingly, when we doubled the initial concentration of actin monomers to 6 μM, the maximum comet length was 130 μm, whereas it was only 60 μm for an initial actin monomer concentration of 3 μM (Figure 1D, 1E). This demonstrates that in microwells, actin comet size is determined by the initial pool of monomers and that this pool is rapidly consumed over time.

Another argument for this limited number of components in microwells comes for the analysis of fluorescence profiles of the comet tails (Figure 1G). Indeed, actin fluorescence is constant over the entire tail of the comet in flow chamber (Figure 1G), but decreases in the region of the comet near the beads in microwells (Figure 1G). As new actin filaments are nucleated at the surface of the bead, this result confirms that the amount of polymerized actin in the wells decreases over time. Therefore, we sought conditions under which the actin monomer pool is maintained overtime to improve the lifetime of actin assembly and bead motility.

### Sustained actin assembly in cell-sized compartment requires disassembly and recycling

We first added the protein ADF/cofilin to our initial mix because it is known to be a major actor of actin dynamics in cells (Lappalainen & Drubin, 1997; Dawe *et al*, 2003; Vitriol *et al*, 2013) and in reconstituted systems (Loisel *et al*, 1999; Suarez *et al*, 2011; Wioland *et al*, 2017). Actin comet tail disassembly was induced by the addition of 200 nM of ADF/cofilin to our initial mixture. The choice of the ADF/cofilin concentration was based on our previous work on branched actin dynamics (Manhart *et al*, 2019) and the concentration dependence of ADF/cofilin activity (Andrianantoandro & Pollard, 2006). In these conditions, actin comets do not just assemble for a limited time (Figure 2A, Supplemental Figure 2A, Movie 3, Assembly conditions) but assemble and disassemble (Figure 2B, Supplemental Figure 2B, Movie 4, Disassembly conditions). The disassembly step following the addition of ADF/cofilin allowed the bead to move for ∼24 hours (right panel in Figure 2B, Figure 2D and movie 2 middle column) instead of only ∼3 hours in the absence of ADF/cofilin (right panel in Figure 2A, Figure 2D and movie 2 first column). In consequence, the half-life of bead motility was increased in disassembly conditions compared to assembly conditions (0.6 hours in Assembly conditions *vs*. 2 hours in Disassembly conditions, Figure 2E). However, even in the presence of ADF/cofilin, the bead velocity decreased rapidly within the first 5 hours of the experiment (Figure 2 F and G). Since the pool of monomers is limited in our conditions, one possibility for the non-stability of the system is its inefficiency to recycle actin subunits or fragments generated by ADF/cofilin-induced disassembly to recharge the pool of actin monomers bound to ATP.

In order to improve the recycling steps of the actin turnover, we added cyclase-associated protein (CAP), which is well known to be important for actin dynamics in cells (Iwanski *et al*, 2021; Schneider *et al*, 2021; Rust *et al*, 2020). In addition, CAP is known to have a synergic effect with ADF/cofilin during actin disassembly, as demonstrated in cells (Bertling *et al*, 2004) and in *in vitro* (Normoyle & Brieher, 2012; Chaudhry *et al*, 2013; Kotila *et al*, 2019; Shekhar *et al*, 2019). CAP also catalyzes nucleotide exchange more efficiently than the profilin (Moriyama & Yahara, 2002; Chaudhry *et al*, 2010), therefore potentially playing a key role during actin monomer recycling. Addition of 400 nM of CAP, 2 excess molar ratio over ADF/cofilin ((Chaudhry *et al*, 2014; Kotila *et al*, 2019), Figure 2C, Supplemental Figure 2C, Movie 5, Recycling conditions) further enhanced the half-life of bead motility (8 hours in Recycling conditions *vs*. 2 hours in Disassembly conditions, Figure 2D, 2E and movie 2 right column). Importantly, in most cases, bead velocity and comet surface area became nearly constant for 4 hours (Figure 2F, 2G), as expected for a dynamic steady state. These results show that we succeeded in reconstituting a dynamic steady state in a cell-sized compartment with an average lifetime of 4 hours. However, this lifetime can be limited by additional factors that we will address in the following parts of this study.

### The actin monomer pool is recycled several times to maintain actin assembly in cell-sized compartment

We sought to determine the amount of actin assembled in the comet as a function of time to assess the number of times the assembly/disassembly/recycling cycle was performed during sustained bead motility. First, in assembly conditions, we determined the amount of actin in the comet tails over time, by measuring the total fluorescence intensity of actin in the microwells and in the comet tail (Figure 3A). Given the critical concentration of ATP-actin (0.1 μM), full polymerization should theoretically result in ∼ 96 % actin in comet tails. We estimate that in our condition 57 % of the actin initially introduced in the microwell assembles into the comet tail (the rest being likely polymerized in the bulk) (Figure 3A). Quantitative estimates considering actin concentration and comet length confirm this value. (see Materials and Methods).

**Figure 3.**
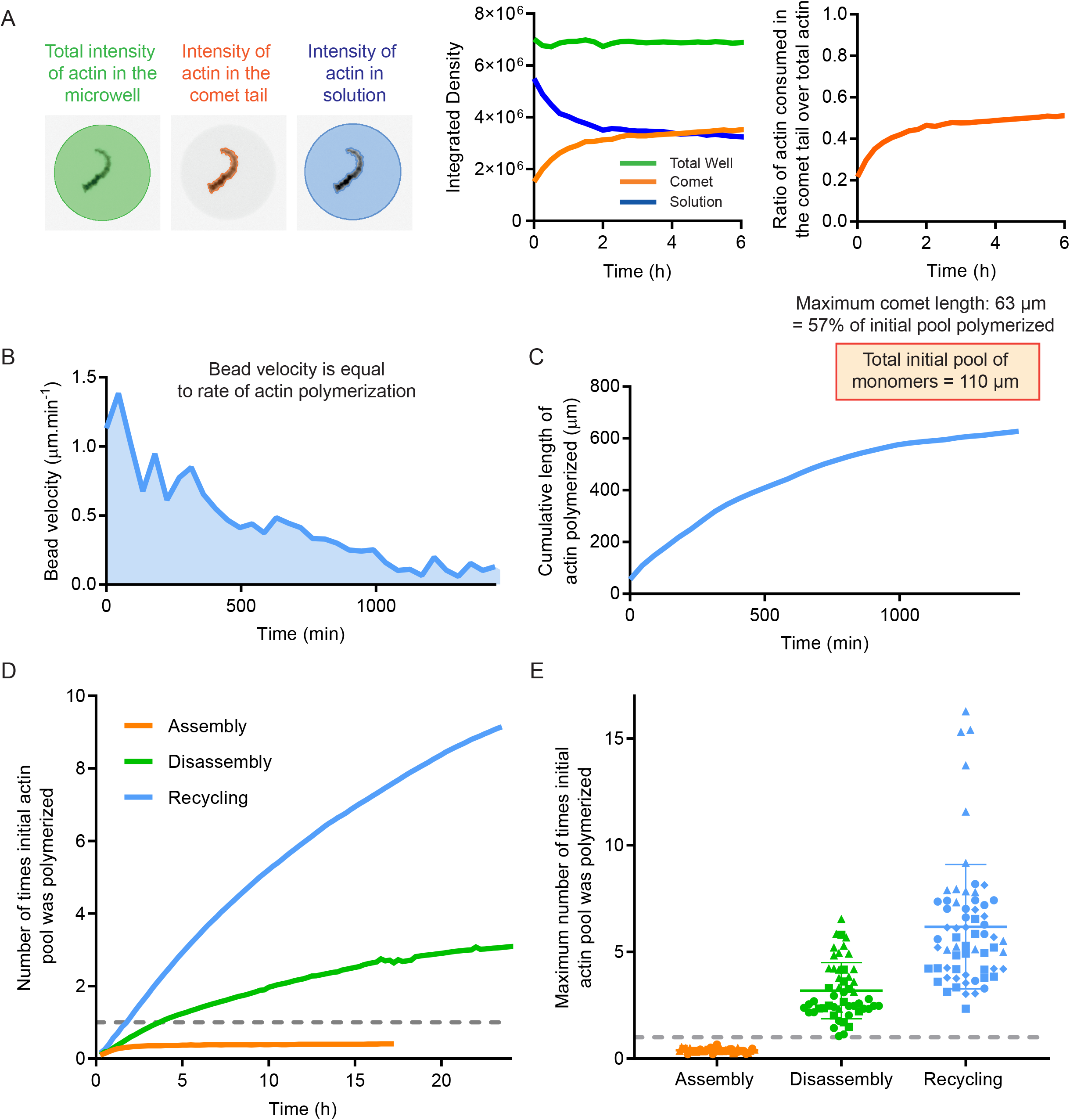
Actin monomers are recycled several times during sustained actin turnover in cell-sized compartment. **A**. Left: Estimation of the total actin amount in microwell and in the comet tail. Middle: Actin integrated density as a function of time for comet tail, solution and total well. Right: Ratio of actin consumed in comet tail versus total actin as a function of time. **B**. Example of bead velocity as a function of time in recycling conditions. The integral of this curve (or area under curve) represents the length of actin polymerized in the comets during the time of the experiment. **C**. Example of cumulative length of actin polymerized as a function of time. **D**. Mean number of times initial actin amount was polymerized as a function of time for one dataset per condition. **E**. Maximum number of times the initial pool of actin monomers was polymerized in the microwell for the different conditions for actin-based motility. The grey dashed line represents 1 cycle, which is equivalent to 3 μM, the initial concentration of actin introduced in the microwell. Assembly: N = 3, n= 38 comet tails. Disassembly: N = 3, n = 52 comet tails. Recycling: N = 4, n= 65 comet tails. 1 symbol per independent dataset.

In disassembly and recycling conditions, we computed the integrals of the bead velocity curves as a function of time (Figure 3B) in order to estimate the quantity of actin polymerized in the comet tail over time. Indeed, by computing the integral of the velocity curve, we obtained the cumulative length of actin polymerized as a function of time (Figure 3C). In order to convert the actin length into a quantity of actin polymerized (in μM), we used the fact that under assembly conditions, 57% of the actin initially introduced in the microwell assembles into the comet tail. As the mean actin comet length is 63 μm, the total initial pool of actin monomers represents ∼110 μm of comet tail. We used this factor to convert the length of actin polymerized in a quantity of actin polymerized. From those measurements, we were able to estimate the consumption of actin monomers by the system. In recycling conditions, the whole system consumed initially about 2 μM of actin monomer per hour whereas it was slower in assembly and disassembly conditions (Figure 3D). In disassembly conditions, the initial pool of actin monomers was assembled in the comet on average 3 times (Figure 3E), demonstrating that the presence of ADF/cofilin and profilin enables recycling of ADP-actin subunits after disassembly as suggested before (Blanchoin & Pollard, 1998). In recycling conditions, the addition of CAP further improved the ability of the system to reuse the actin monomer pool in multiple cycles of actin turnover. Indeed, in presence of CAP, the initial pool of actin monomers was assembled on average 6 times (and up to 17 times, Figure 3E).

The disassembly rate of the actin network was measured experimentally by tracking the fluorescence of actin defects in the comets, and examining their fluorescence decay over time (Supplemental Figure 3A, B and C). Interestingly, under disassembly conditions, the mean disassembly time of the comet was 55 minutes whereas it was estimated to be 22 minutes on average under recycling conditions (Supplemental Figure 3D). When the disassembly time was “normalized” to the comet length, we were able to estimate a rate of network disassembly, which in both disassembly and recycling conditions was approximately 1.2 μm.min^-1^ matching the rate of actin assembly (Supplemental Figure 3E).

Although our system approaches on average a dynamic steady state for 4–hours under recycling conditions, the bead velocity decreases over time, and bead motility eventually stops (Figure 2D). This result suggests the presence of a limiting factor affecting the lifetime of our system. Since actin assembly consumes one ATP each time an actin monomer adds to the comet tail, we tested whether ATP depletion explains the decreasing velocity.

### ATP is necessary to maintain the dynamic steady state, but is not the limiting factor

We varied ATP concentration from 7 μM (coming only from the actin introduced in the reaction mix) to 3 mM (concentration used in the previous parts of the study) under assembly, disassembly and recycling conditions (Figure 4, Movie 6 and Supplemental Figure 4). Interestingly, under assembly conditions, the initial concentration of ATP has minimal influence on the kinetics of comet growth (Supplemental Figure 4A) and the comet area and bead velocity were similar for both ATP concentrations (Supplemental Figure 4B, C). In contrast, ATP concentration has an effect in the presence of ADF/cofilin after the initial 2 hours. At 3 mM ATP, comets started to disassemble and the length of the comets decreased while at low ATP concentration, this disassembly was less efficient (Supplemental Figure 4D-F). Of particular interest, the number of polymerization cycles of the initial monomer pool was independent of ATP concentration under assembly conditions (Supplemental Figure 4G) but increased significantly with higher ATP concentration under disassembly conditions (Supplemental Figure 4G).

**Figure 4.**
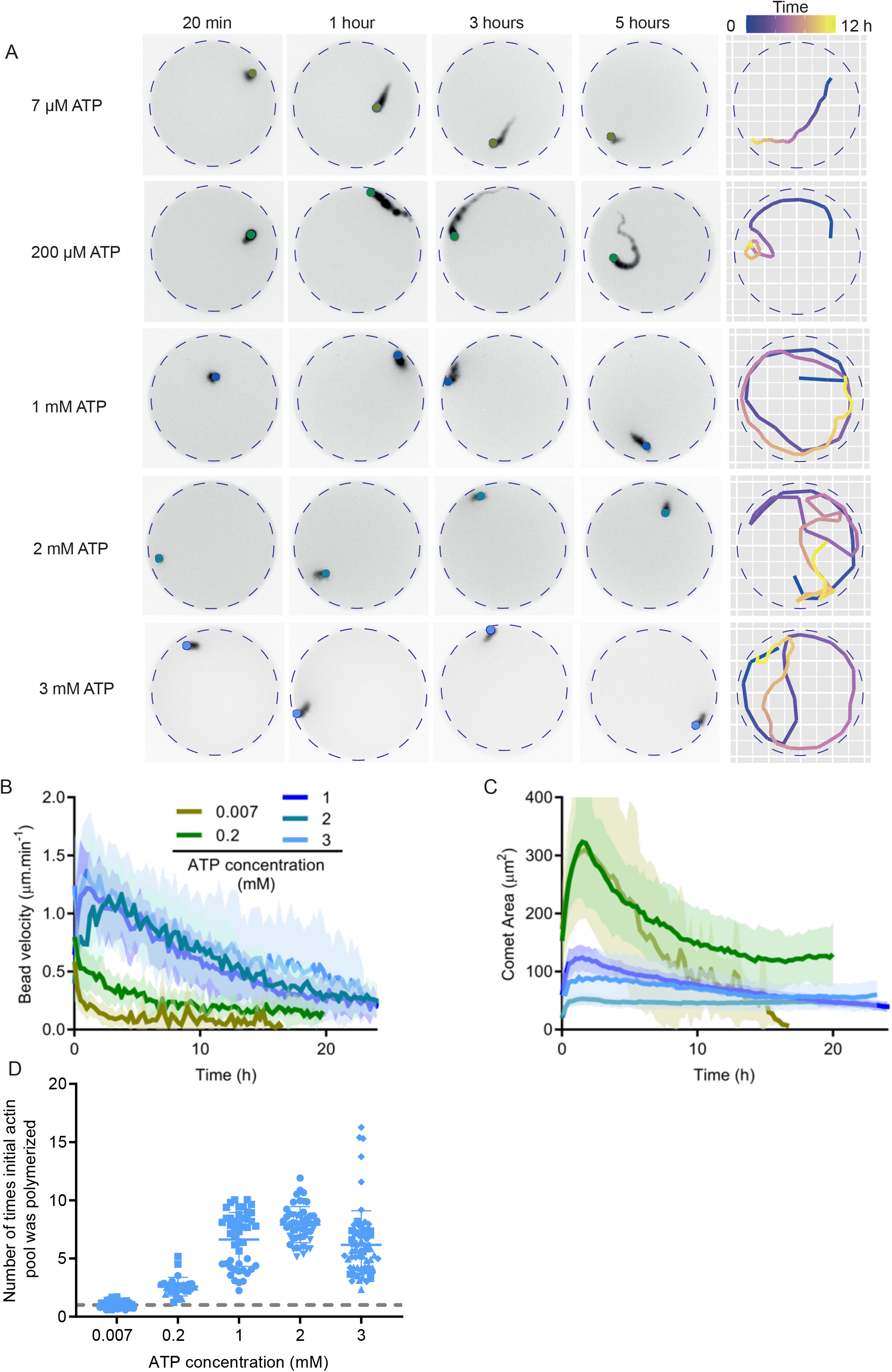
ATP is necessary for sustained actin turnover. **A**. Left: Snapshots of actin comet tails in recycling conditions with the indicated ATP concentrations. Right: Tracking of the comet shown in the snapshots. Time is encoded in color. **B**. Bead velocity as a function of time for one dataset per condition. Mean and standard deviation are represented. [ATP] = 0.007 mM: n = 12 comet tails. [ATP] = 0.2 mM: n = 16 comet tails. [ATP] = 1 mM: n = 30 comet tails. [ATP] = 2 mM: n = 31 comet tails. [ATP] = 3 mM: n = 16 comet tails. **C**. Comet area as a function of time for one dataset per condition. Mean and standard deviation are represented. **D**. Number of times initial actin quantity was polymerized in the microwell for various concentrations of ATP. The grey dashed line represents 1 cycle which is equivalent to 3 μM, the initial concentration of actin introduced in the microwell. [ATP] = 0.007 mM: N=2, n = 35 comet tails. [ATP] = 0.2 mM: N = 3, n = 30 comet tails. [ATP] = 1 mM: N = 2, n = 47 comet tails. [ATP] = 2 mM: N = 1, n = 3l comet tails. [ATP] = 3 mM: N = 4, n = 65 comet tails. Biochemical conditions: 4.5 μm polystyrene beads coated with 400 nM Strep-SNAP-WA; 3 μM actin, 6 μM profilin, 90 nM Arp2/3, l5 nM Capping protein, 200 nM cofilin, 400 nM Cyclase Associated Protein (CAP) and ATP concentration as indicated.

We then varied the ATP concentrations under recycling conditions (Figure 4A, Movie 6), adding several intermediate concentrations (200 μM, 1 mM and 2 mM) to the 7 μM and 3 mM conditions used previously. At low ATP concentrations (7 and 200 μM), comet dynamics resemble those of disassembly conditions, with an initial increase in comet area followed by a progressive decrease (Figure 4C), a rapid decrease in bead velocity over time (Figure 4C), and almost complete comet disassembly after 8 hours (Figure 4A, C, Movie 6). At high ATP concentrations (1, 2 and 3 mM), there is a dramatic increase in both bead velocity and comet area, although both decrease progressively over 20 hours (Figure 4A, B and C). Estimation of the number of times the initial actin monomer pool was consumed for the different ATP concentrations revealed that 1 mM ATP is already a saturating ATP concentration (Figure 4D).

These results demonstrate that recycling generates faster and long-lived systems but that are more sensitive to energy input. Since energy was not the limiting factor to maintain the steady state in our experimental conditions, we hypothesized that another component of our system may have degraded during the experiment. To test this hypothesis, we decided to evaluate the aging of the different components of the system in our experiment.

### Actin aging limits the lifetime of actin assembly

The microwells are enclosed between a slide and a coverslip, implying that their content cannot be modified during the experiment. Therefore, addressing protein aging in microwells was technically challenging. We decided to compare under recycling conditions the slow decrease in bead velocity in the flow chamber (unlimited number of components) and in the microwells (limited number of components). We observed that the rate of decrease of 0.1 h^−1^ was similar in the two conditions (Supplemental Figure 5A). We therefore decided to investigate possible mechanism responsible for this decrease in the flow chamber.

**Figure 5.**
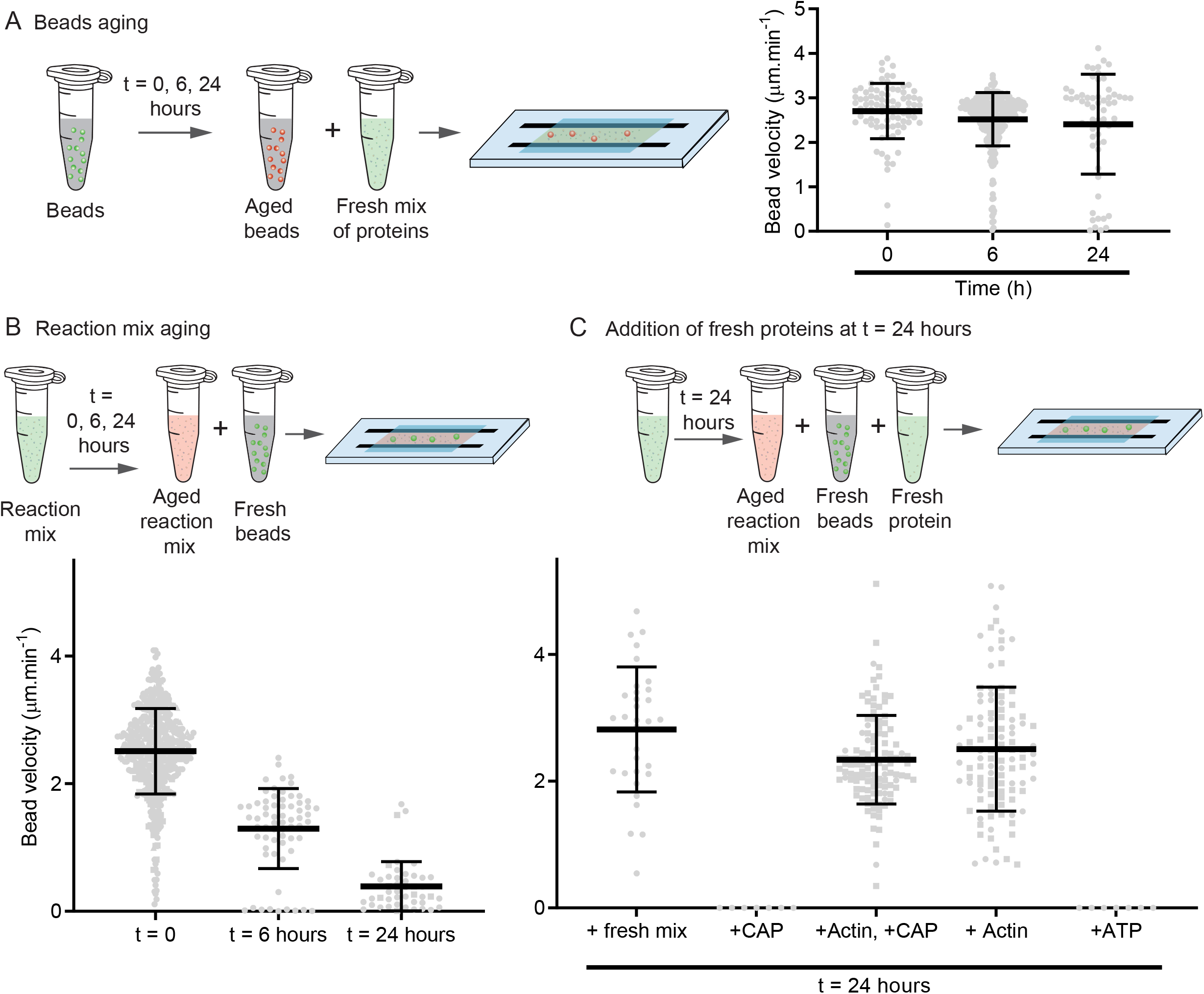
Aging of components during the actin dynamic steady state. **A**. Test of beads aging in motility buffer. Beads coated with 400 nM of Strep-SNAP-WA were left on the bench and reintroduced into a new protein mixture at different elapsed times (0, 6 and 24 hours). Bead velocity was estimated for those different times. Composition of the protein mix: 3 μM actin, 6 μM profilin, 90 nM Arp2/3, 15 nM Capping protein, 200 nM cofilin, 400 nM Cyclase Associated Protein (CAP). N = 1, n = 87 comets for t = 0, n = 292 comets for t = 6 hours, n = 60 comets for t = 24 hours. **B, C**. Test of reaction mix aging without beads. Reaction mix was prepared without beads and left on the bench at room temperature. Then at different time intervals, fresh beads were added to the mix. In some conditions (right part of the graph), fresh proteins were added with the beads. Velocity of fresh beads was estimated for those different times. Composition of the reaction mix: 3 μM actin, 6 μM profilin, 90 nM Arp2/3, 15 nM Capping protein, 200 nM cofilin, 400 nM Cyclase Associated Protein (CAP). t = 0: N = 4, n = 494 comets. t = 6 hours: N = 1, n = 65 comets. t = 24 hours: N = 2, n = 46 comets. t = 24 hours + fresh mix: N = 1, n = 32 comets. t = 24 hours + actin: N = 2, n = l08 comets. t = 24 hours + actin + CAP: N = 2, n = 115 comets.

We first examined possible aging of the bead component, that is, whether the NPF proteins on the bead could become unstable or detach from the bead over time (Figure 5A). We prepared NPF-coated beads, age them at room temperature for 0, 6 or 24 hours before adding them to a fresh motility mixture (Figure 5A, Movie 7). The bead velocity was similar at these different time intervals (Figure 5A), suggesting that bead aging did not have a significant impact on the lifetime of our system.

We next tested the lifetime of the reaction mixture by allowing the reaction to age before adding fresh beads (Figure 5B). Bead velocity gradually decreased with the aging time of the reaction mixture (Figure 5B, Movie 8), losing all motility after 24 hours. These experiments demonstrate that protein aging in the reaction mixture is a limiting factor for the lifetime of the system. Moreover, the addition of a “fresh” mixture of proteins to the aged reaction allowed the beads to restore their initial velocity (Figure 5C, Movie 9).

To identify the proteins that were aging in our system, we performed high-speed centrifugation assays (Supplemental Figure 5B, Materials and methods). We introduced each protein of our assay separately in the motility buffer and we left it at room temperature for aging. We then looked at the amount of protein in the high-speed pellet at t = 0 and t = 24 hours. From this assay, we identified that most proteins of our assay were stable in the supernatant overtime except for the CAP protein, which was found in a higher amount in the pellet after 24 hours (Supplemental Figure 5B). The only protein that we could not test with the pelleting assay was the actin (because it would go to the pellet as soon as it forms filaments).

We therefore tested the effect of adding CAP and/or actin after 24 hours of aging in our motility assay (Figure 5C, Movie 9). When CAP was added to an aged mix after 24 hours, the beads did not move. Interestingly when actin or actin with CAP were introduced after 24 hours, the beads resumed movement with a velocity similar to the one observed at the initial time of the experiment. As a control, we showed that it was not the small amount of ATP brought by actin that restarted the system by adding 3 mM ATP after 24 hours to our system. Experiments where we aged CAP individually in a test tube demonstrated bead velocity was similar with fresh CAP and with CAP aged for 24 hours (Supplemental Figure 5C). Therefore, CAP does not age significantly enough in our experiment to reduce bead motility over time. On the contrary, experiments where actin was aged individually in a test tube demonstrated that bead velocity was lower for actin aged for 24 hours than for fresh actin (Supplemental Figure 5D). We thus conclude that actin monomers are the limiting factor in the system and their aging cause motility to decrease over time.

## Discussion

Our results established the relative contribution of the three-step cycle (assembly, disassembly, recycling) on actin bead motility in the presence of a limited pool of actin monomers. This allows us to propose a quantitative model of the control of actin turnover lifetime in a cell-sized compartment for different scenarios.

Under assembly conditions, at first, actin tail grows rapidly and consumes the actin monomer pool (Figure 6A, Early Assembly). Later, disassembly is too slow to compensate for the rapid monomer consumption and growth stops (Figure 6A, Assembly Late). This demonstrates that tread milling limited by the rate of depolymerization at the pointed ends of the filament cannot account for rapid actin turnover (Pollard & Borisy, 2003; Miyoshi & Watanabe, 2013; Blanchoin *et al*, 2014). A simple kinetic model (see materials and methods) considering the volume of the microwells, the concentration of the actin pool and the rate of polymerization on the surface of the beads, accurately predicts the variation of the comet growth speed as a function of time. The model also predicts accurately the comet length as a function of the initial pool of actin monomers (Figure 6B).

**Figure 6.**
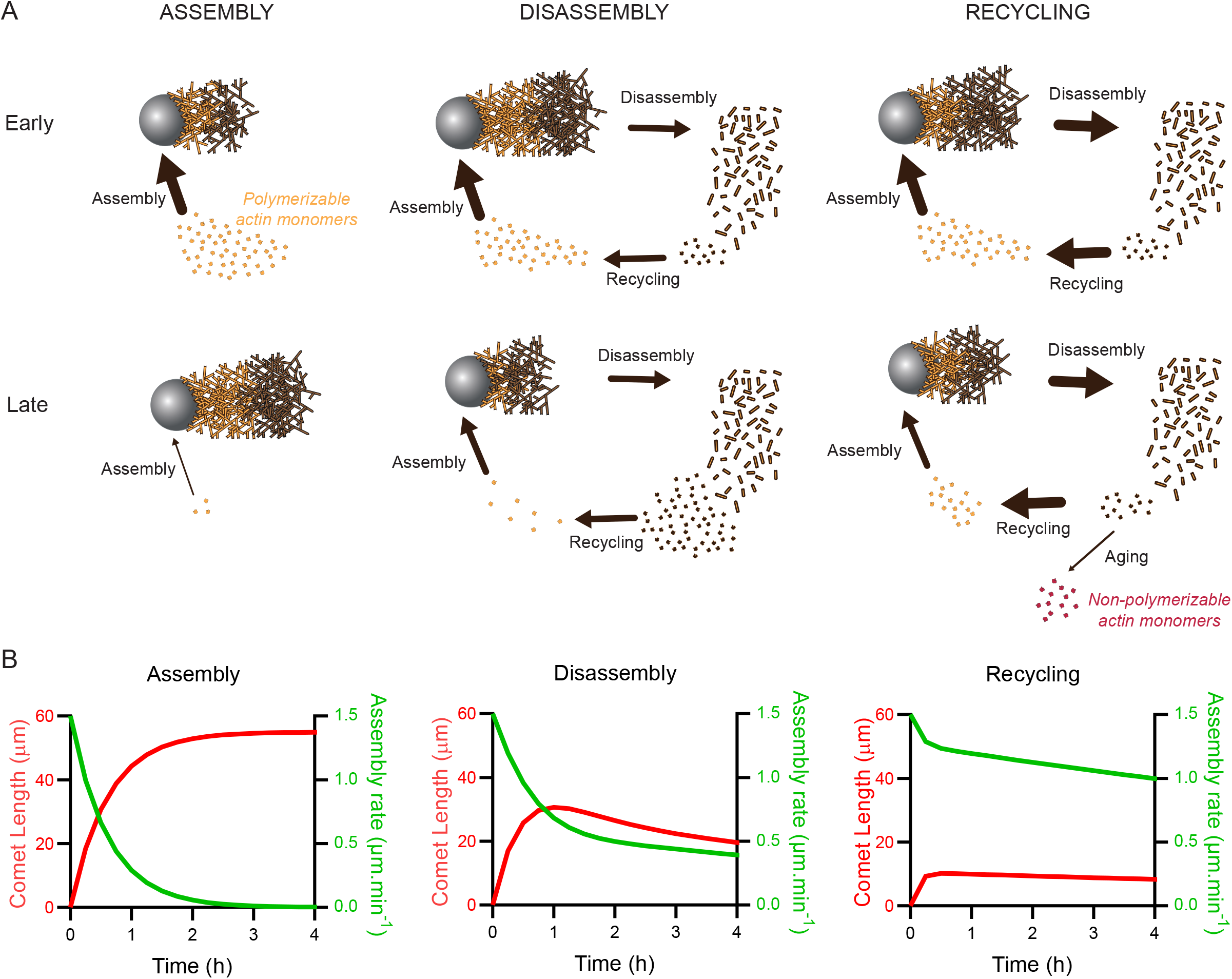
Quantitative model of the control of actin turnover lifetime in a cell-sized compartment. **A**. Summary scheme of the different conditions reconstituted in this study and the fluxes associated. The size of arrows scales with the value of the associated flux. **B**. Model prediction of the assembly rate and comet length as a function of time in assembly, disassembly and recycling conditions.

Under disassembly conditions, the initial rate of actin assembly is faster than disassembly (Figure 6A, Disassembly early). At early stage, the comet grows and the monomer pool decreases. As a result, the actin assembly rate decreases and at a given time, corresponds to the disassembly rate. The comet then reaches a stable length over time. The actin network dynamic steady state is then limited by the recycling step necessary to maintain the pool of actin monomer (Figure 6A, Disassembly late). Interestingly, this system exhibits a feedback loop between assembly and disassembly via the monomer pool that allows the system to adapt to variations in operating rate over time (Figure 6B). Then, a dynamic steady state of the actin network can be reached only in rare cases, when the initial assembly rate precisely matches the disassembly rate (Akin & Mullins, 2008; Manhart *et al*, 2019). Implementation in our model of the disassembly and recycling in presence of ADF/cofilin shows their impact on the length and growth speed of actin comets (Figure 6B).

Under recycling conditions, the rate of disassembly and recycling increases (see materials and methods for an estimate of these rates in disassembly and recycling conditions). The comet reaches a smaller stable length (Figure 6A, Recycling early). Fast recycling maintains the pool of actin monomer and increases the system lifetime (Figure 6A, Recycling late). The rate of actin network turnover in this condition (approximately 1 μm.min^-1^, Figure 6B Recycling) is very close to the rate measured *in vivo* for the lamellipodium (Wang, 1985; Theriot & Mitchison, 1991; Watanabe & Mitchison, 2002), demonstrating that we were able to reconstruct actin turnover at physiological rates in a cell-sized compartment.

Fast recycling step was achieved by the protein CAP, which has emerged as a key regulator of actin dynamics *in vivo* (Rust *et al*, 2020). CAP works synergistically with ADF/cofilin to depolymerize actin filaments and subsequently catalyzes nucleotide exchange on the resulting monomers (Kotila *et al*, 2018, 2019; Shekhar *et al*, 2019). Our experiments provide evidence that, similar to cells, both acceleration of actin filament end depolymerization and nucleotide exchange on monomeric actin by CAP are important to reconstitute actin turnover at physiological rates.

Our work also emphasizes the need for a high recycling rate to maintain the pool of actin monomers necessary to participate in the assembly reaction. Most cells possess the complete recycling machinery that is composed of profilin and CAP (Paavilainen *et al*, 2004). However, in the absence of efficient monomer recycling, cells can use alternative strategies such as assembly by oligomer annealing which has the advantage of being energetically more favorable (Okreglak & Drubin, 2010; Smith *et al*, 2013). Another possibility to overcome fast recycling rate is to maintain a large pool of non-polymerizable monomers (most likely bound to thymosin-ß4) that acts as a buffer allowing a temporary mismatch between assembly and recycling (Vitriol *et al*, 2015; Raz-Ben Aroush *et al*, 2017).

Actin turnover and the dynamic steady state of the network are intimately linked to energy consumption. Our results show that while assembly is independent of free ATP concentration, as long as an ATP is bound to an actin monomer (Blanchoin & Pollard, 1999), disassembly and recycling are very sensitive to ATP concentration. At low ATP concentration, recycling, which involves the exchange of the nucleotide bound to the actin monomer (Blanchoin & Pollard, 1998), is the limiting step in actin turnover. However, at high ATP concentration, recycling is almost instantaneous and disassembly becomes the limiting step. Interestingly, above 1 mM free ATP, the energy supply is not the factor defining the lifetime of our dynamic system. Since the concentration of ATP in a physiological context is well above 1 mM (Greiner & Glonek, 2021), our findings reinforced the notion that cells contained excessively high ATP concentration, compared to the concentration necessary to maintain actin organization at a dynamic steady state.

In addition to recycling, aging of actin monomers limits the lifetime of the system. Lifetime of *in vitro* systems has been overlooked in previous studies. Very few studies mentioned the lifetime of bead motility in unlimited volumes (Marchand *et al*, 1995; Akin & Mullins, 2008; Lacayo *et al*, 2012). In our case, actin monomer aging is most likely occurring due to the presence of reactive oxygen species (ROS) that induce thiol modifications of cysteine residues. Cys374 on actin is highly reactive and probably the most redox-sensitive cysteine (Lassing *et al*, 2007). Cysteine oxidation can cause the formation of intra- or inter-molecular disulfide bridges (Farah *et al*, 2011; Balta *et al*, 2021; Rouyere *et al*, 2022) that impairs actin polymerization (DalleDonne *et al*, 1995, 1999). As one of the intermediates during nucleotide exchange is nucleotide-free actin which is very unstable unless stabilized by sucrose (De La Cruz & Pollard, 1995), it can be thought that the repetition of cycles during turnover may induce its degradation. It is also possible that during these successive cycles, actin becomes more sensitive to ROS. Cells may have overcome these issues with systems of chaperones (Dalle-Donne *et al*, 2001; Wettstein *et al*, 2012; Grantham, 2020), by having an effective system of protein synthesis and degradation to maintain a pool of polymerizable actin monomers (Olson & Nordheim, 2010; Vedula *et al*, 2021) or by limiting oxidative stress (Rouyere *et al*, 2022).

To achieve structural stability and dynamics in cellular organization, the importance of balancing the supply and demand of building blocks in real time is crucial. Since cellular building blocks are often limited, recycling seems to be essential to keep this balancing mechanism under control and avoid a mismatch between supply and demand that would alter structural stability. Our system offers new opportunities to study the essential role of recycling in the balancing mechanism as a general principle for the dynamic steady state of intracellular organization. With this work, reconstituted systems were pushed to the limits that are key to cellular life: recycling and energy production to feed active assembly/disassembly as well as component self-renewal to limit aging.

## Figure legends

**Supplemental Figure 1.**
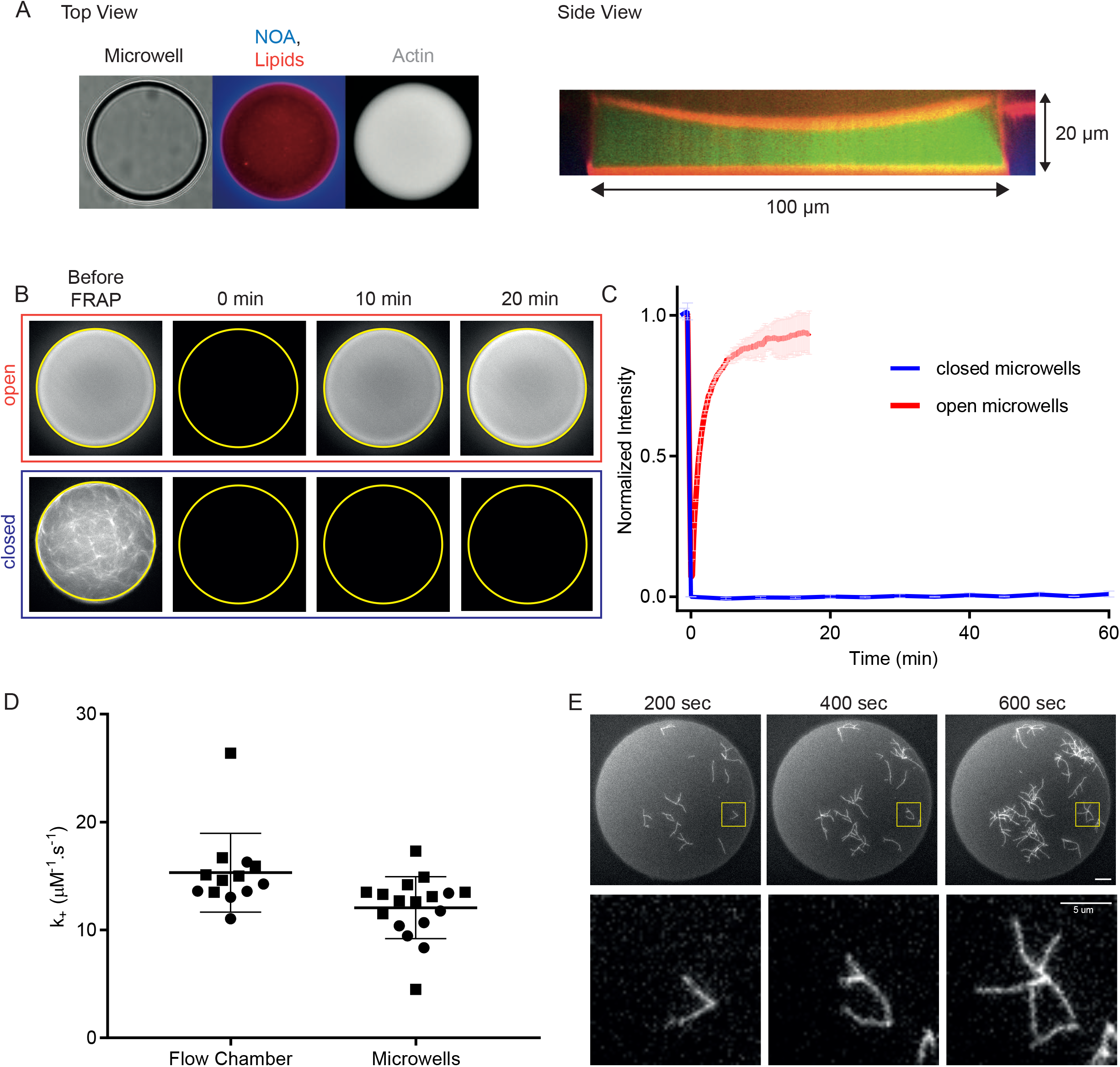
Microwells are closed environments that preserve key parameters for actin assembly. **A**. Top and side views of microwells used in this study. **B**. Snapshots of FRAP experiment in open (Top) and closed (Bottom) microwells. **C**. Quantification of FRAP experiment in open and closed microwell. Closed wells: N = 3, n = 5 microwells. Open wells: N = 1, n = 2 microwells. **D**. Quantification of the association rate constant of actin filament assembly at the barbed ends in flow chamber and in microwells. Biochemical conditions: [actin] = 0.8 μM. [profilin] = 2.4 μM. N = 2, n = 13 filaments for the flow chamber and n = 17 filaments for the microwells. **E**. Visualization of actin branched network formation in closed microwells (full well and zoom). Biochemical conditions: [actin] = 1 μM. [profilin] = 3 μM. [WA] = 50 nM. [Arp2/3 complex] = 25 nM.

**Supplemental Figure 2.**
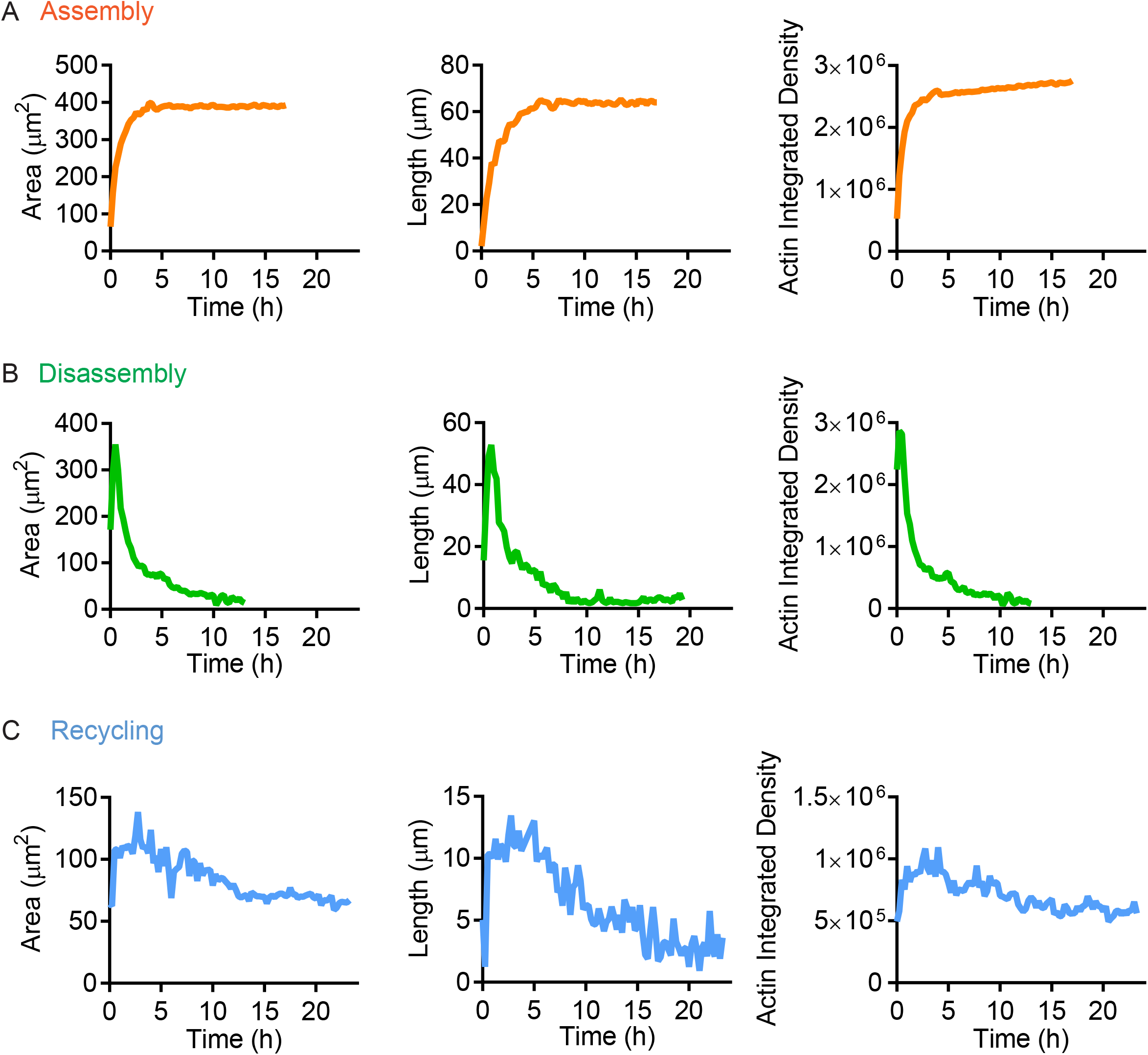
Quantitative analysis of actin within the comets in assembly, disassembly and recycling conditions. **A**. Estimation of Length, Area and Actin integrated density of the comet shown in Figure 2 for Assembly conditions. **B**. Estimation of Length, Area and Actin integrated density of the comet shown in Figure 2 for Disassembly conditions. **C**. Estimation of Length, Area and Actin integrated density of the comet shown in Figure 2 for Recycling conditions.

**Supplemental Figure 3.**
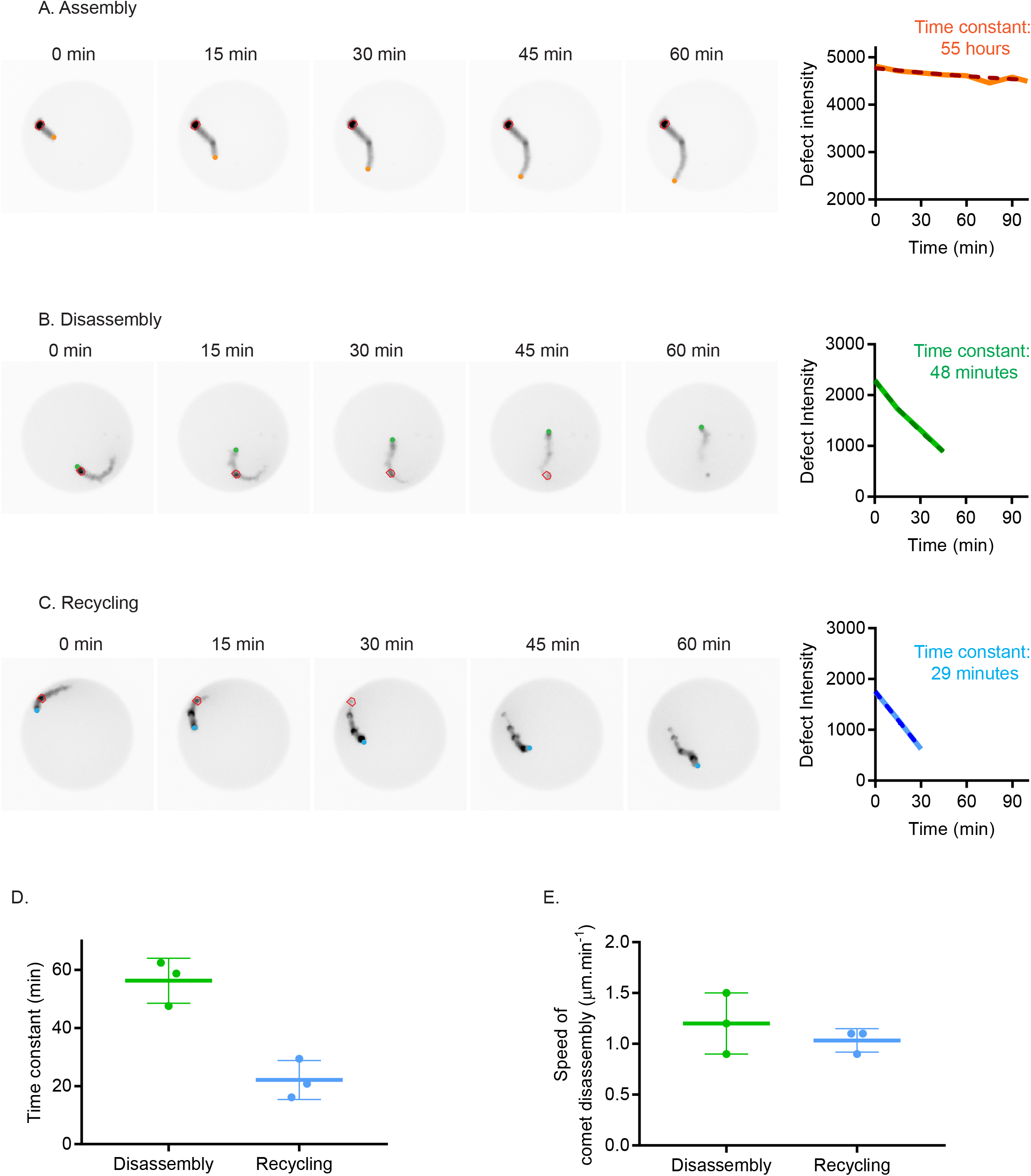
Determination of the rate of disassembly of the actin comets in assembly, disassembly and recycling conditions. **A**. Left: Snapshots of an actin comet tail assembled in Assembly conditions in a microwell. The bead is in orange and the tracked defect in red. Right: Defect fluorescence intensity as a function of time (solid line) and exponential fit (dashed line). Time constant is estimated from the exponential fit. **B**. Left: Snapshots of an actin comet tail assembled in Disassembly conditions in a microwell. The bead is in green and the tracked defect in red. Right: Defect fluorescence intensity as a function of time (solid line) and exponential fit (dashed line). Time constant is estimated from the exponential fit. **C**. Left: Snapshots of an actin comet tail assembled in Recycling conditions in a microwell. The bead is in blue and the tracked defect in red. Right: Defect fluorescence intensity as a function of time (solid line) and exponential fit (dashed line). Time constant is estimated from the exponential fit. **D**. Comet disassembly time in disassembly (N = 2, n = 3 comets) and recycling (N = 2, n = 3 comets) conditions. **E**. Speed of comet disassembly in disassembly and recycling conditions.

**Supplemental Figure 4.**
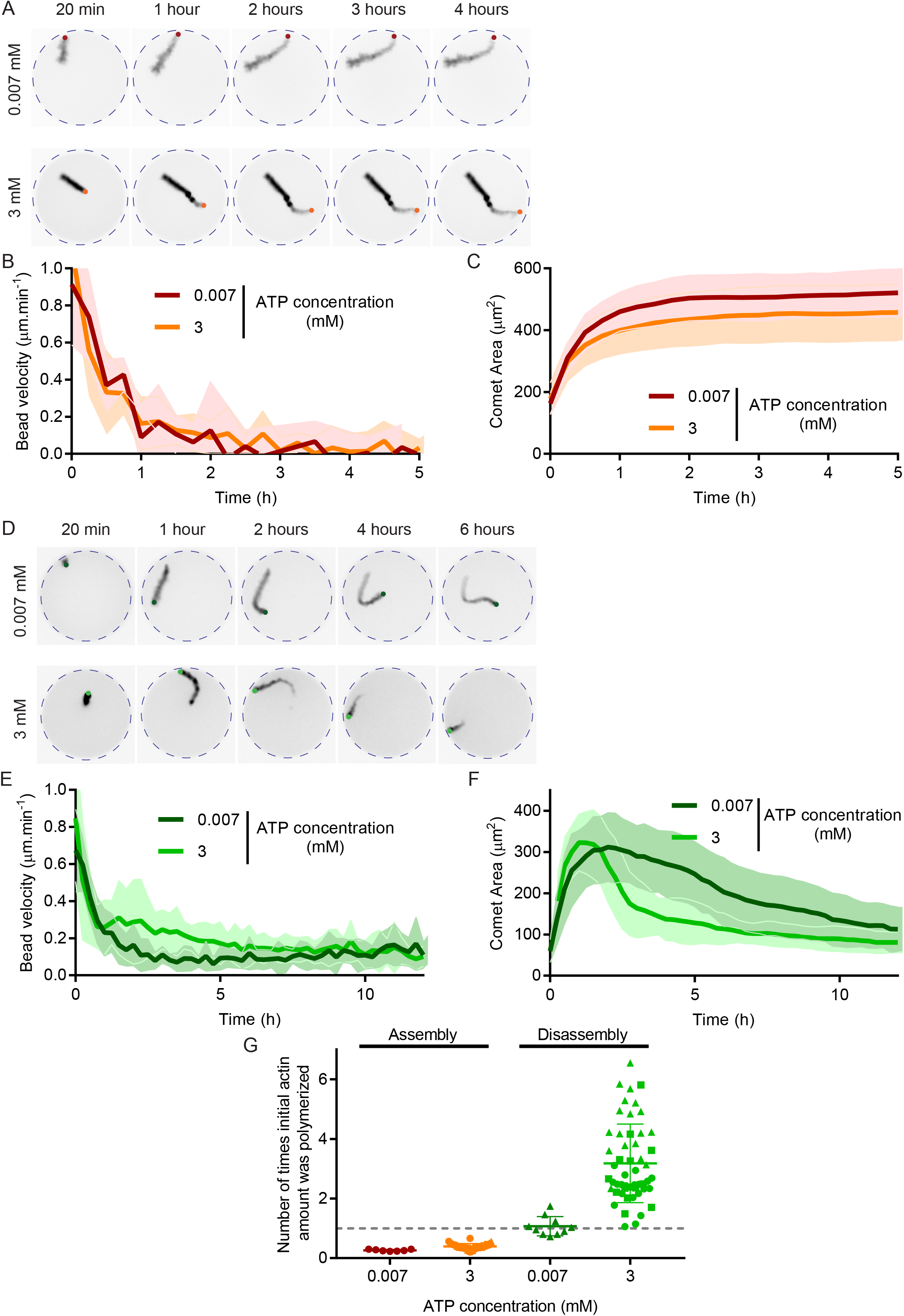
Effect of ATP concentration on actin-based motility in assembly and disassembly conditions. **A**. Snapshots of actin comet tails in assembly conditions with low ([ATP] = 0.007 mM) or high ([ATP] = 3 mM) ATP concentrations. **B**. Quantification of bead velocity for one dataset per condition in assembly conditions for low ([ATP] = 0.007 μM) or high ([ATP] = 3 mM) ATP concentrations. [ATP] = 0.007 μM: 7 comet tails. [ATP] = 3 mM: 11 comet tails. **C**. Quantification of comet area for one dataset per condition in Assembly conditions. **D**. Snapshots of actin comet tails in disassembly conditions with low ([ATP] = 0.007 mM) or high ([ATP] = 3 mM) ATP concentrations. **E**. Quantification of bead velocity for one dataset per condition in disassembly conditions for low ([ATP] = 0.007 μM) or high ([ATP] = 3 mM) ATP concentrations. [ATP] = 0.007 mM: 10 comet tails. [ATP] = 3 mM: 16 comet tails. **F**. Quantification of comet area for one dataset per condition. **G**. Number of times initial actin quantity was polymerized in the microwell for various concentrations of ATP in assembly or disassembly conditions. The grey dashed line represents 1 cycle which is equivalent to 3 μM, the initial concentration of actin introduced in the microwell. Assembly, [ATP] = 0.007 mM: N = 1, n = 7 comet tails. Assembly, [ATP] = 3 mM: N = 3, n = 38 comet tails. Disassembly, [ATP] = 0.007 mM: N = 1, n = 10 comet tails. Disassembly, [ATP] = 3 mM: N = 3, n = 52 comet tails. 1 symbol per independent dataset. Biochemical conditions: 4 μm polystyrene beads coated with 400 nM WA; 3 μM actin, 6 μM profilin, 90 nM Arp2/3, 15 nM Capping protein (Assembly), 200 nM cofilin (Disassembly).

**Supplemental Figure 5.**
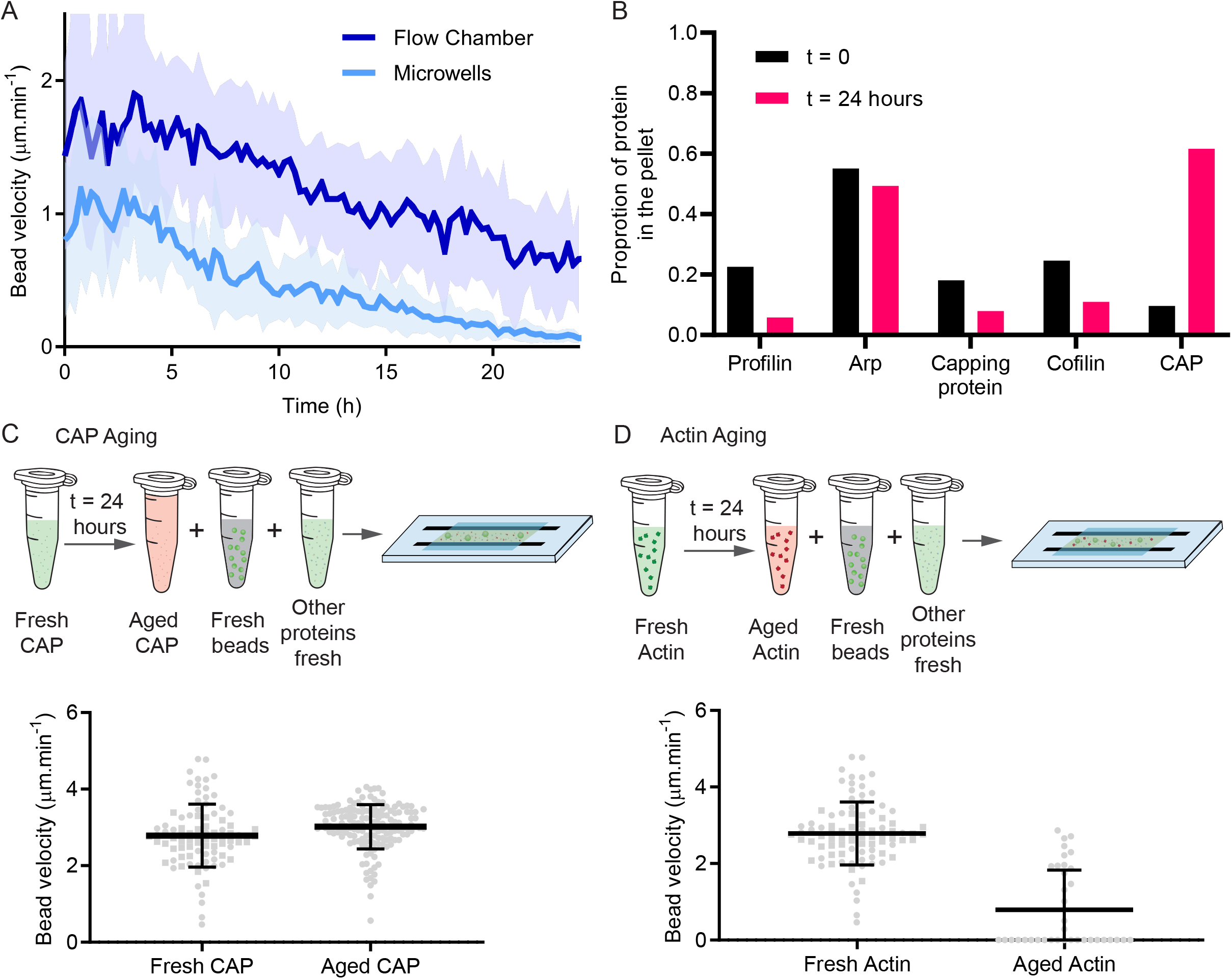
Determination of the aging factor in our biochemical assay. **A**. Comparison of bead velocity in flow chamber and in microwells (one dataset in each condition, n = 37 comet tails in flow chamber; n = 13 comet tails in microwells). **B**. Quantification of the pelleting assay. For each protein, at t = 0 and t = 24 hours, the ratio between the quantity of protein found in the pellet over the total quantity of protein was quantified. **C**. Test of CAP aging. CAP was diluted in motility buffer and left on the bench at room temperature overnight. The morning after, the protein was added to the motility assay. Velocity of beads was estimated in the different conditions. Composition of the reaction mix: 3 μM actin, 6 μM profilin, 90 nM Arp2/3, 15 nM Capping protein, 200 nM cofilin, 400 nM Cyclase Associated Protein (CAP). Fresh CAP: N = 2, n = 89 comets. Aged CAP: N = 1, n= 149 comets. **D**. Test of actin mix aging. Actin mix was prepared in motility buffer and left on the bench at room temperature overnight. The morning after, the mix was added to fresh beads and fresh proteins (except actin). Velocity of beads was estimated in the different conditions. Composition of the reaction mix: 3 μM actin, 6 μM profilin, 90 nM Arp2/3, 15 nM Capping protein, 200 nM cofilin, 400 nM Cyclase Associated Protein (CAP). N = 2, n= 89 comets for fresh actin, n = 37 comets for aged actin.

## Movies Legends

**Movie 1 - Actin comet tail assembly in bulk and in microwells**.

Time lapse imaging of actin comet tail assembly in bulk and in microwells. Data is also shown in Figure 1D. Movie playback is 10 frames per second. Total elapsed time is 6 hours.

**Movie 2 - Contribution of assembly, disassembly and recycling on actin turnover in cell-sized compartment**.

Time lapse imaging of beads in microwells for various biochemical conditions (Assembly, Disassembly, Recycling). Data is also shown in Figure 2. Movie playback is 20 frames per second. Total elapsed time is 23 hours.

**Movie 3 - Examples of actin comet tails assembled in Assembly conditions in microwells**. Nine examples of time lapse imaging of actin comet tail assembly in Assembly conditions. Movie playback is 10 frames per second. Total elapsed time is 15 hours.

**Movie 4 - Examples of actin comet tails assembled in Disassembly conditions in microwells**.

Nine examples of time-lapse imaging of actin comet tail assembly in Disassembly conditions. Movie playback is 10 frames per second. Total elapsed time is 19 hours.

**Movie 5 - Examples of actin comet tails assembled in Recycling conditions in microwells**. Nine examples of time-lapse imaging of actin comet tail assembly in Recycling conditions. Movie playback is 10 frames per second. Total elapsed time is 22 hours.

**Movie 6 - Variation of ATP concentration in Recycling conditions**.

Time lapse imaging of beads in microwells for various ATP concentrations in recycling conditions. Data is also shown in Figure 4. Movie playback is 20 frames per second. Total elapsed time is 20 hours.

**Movie 7 - Evaluation of beads aging**.

Time lapse imaging of beads left in buffer at room temperature nucleating actin comets in recycling conditions at different time points. Data quantification is shown in Figure 5B. Movie playback is 10 frames per second. Total elapsed time is 3 hours.

**Movie 8 - Evaluation of reaction mix aging**.

Time lapse imaging of beads added to reaction mix left at room temperature for aging at different time points. Data quantification is shown in Figure 5C. Movie playback is 10 frames per second. Total elapsed time is 3 hours.

**Movie 9 - Addition of fresh components after 24 hours of aging**.

Time lapse imaging of beads added with fresh components to reaction mix left at room temperature for aging during 24 hours. Data quantification is shown in Figure 5C. Movie playback is 7 frames per second. Total elapsed time is 160 minutes.

## Materials and methods

### Protein Purification

Actin was purified from rabbit skeletal-muscle acetone powder (Spudich & Watt, 1971). Monomeric Ca-ATP-actin was purified by gel-filtration chromatography on Sephacryl S-300 at 4°C in G buffer (2 mM Tris-HCl, pH 8.0, 0.2 mM ATP, 0.1 mM CaCl_2_, 1 mM NaN_3_ and 0.5 mM dithiothreitol (DTT)). Actin was labelled on lysines with Alexa-568(Isambert *et al*, 1995). All experiments were carried out with 5% labelled actin. The Arp2/3 complex was purified from calf thymus according to (Egile *et al*, 1999) with the following modifications: the calf thymus was first mixed in an extraction buffer (20 mM Tris pH 7.5, 25 mM KCl, 1 mM MgCL2, 0.5 mM EDTA, 5% glycerol, 1 mM DTT, 0.2 mM ATP and proteases). Then, it was placed in a 50% ammonium sulfate solution in order to make the proteins precipitate. The pellet was resuspended in extraction buffer and dialyzed overnight.

Human profilin was expressed in BL21 DE3 pLys *Escherichia coli* cells and purified according to (Almo *et al*, 1994). Mouse capping protein was purified according to (Palmgren *et al*, 2001). Yeast cofilin was purified and fluorescently-labelled according to (Suarez *et al*, 2011). The full-length mouse cyclase associated protein 1 (CAP1) was purified in a similar fashion as described in (Kotila *et al*, 2019). To describe briefly, the CAP1 protein was expressed in *E*.*coli* as described earlier, or by using BL21 (DE3) *E*.*coli* cells (Sigma) and expression in LB medium at +l6°C for 30 hours. The bacteria were pelleted and resuspended to buffer A (50 mM Tris pH 7.5, 150 mM NaCl, 25 mM imidazole) and lysed by sonification in the presence of protease inhibitors (200 μg/ml PMSF, 1 μg/ml leupeptin, 1 μg/ml aprotinin, 1 μg/ml pepstatin A, 150 μg/ml benzamidine hydrochloride, all from Sigma-Aldrich). The supernatant, clarified by centrifugation, was then loaded to a 5 mL HisTrap Ni-NTA column coupled to AKTA Pure protein purification system (GE Healthcare). The His-SUMO-tagged CAP1 protein was eluted from the nickel column with an imidazole gradient using buffer A and buffer B (buffer A + 250 mM imidazole), and the main peak fractions were collected and concentrated with Amicon Ultra-15 30 kDa cutoff centrifugal filter device. The His-SUMO-tag was then cleaved from the CAP protein in the presence of SENP2 protease, afterwhich the cleaved protein was subjected to gel filtration runs by using Superose 6 increase 10/300 GL gel filtration column equilibrated in 5 mM HEPES, 100 mM NaCl, 1 mM DTT, 1 μg/ml leupeptin, pH 7.4. Peak fractions from the same elution volume were combined, concentrated as above and snap-frozen with liquid N_2_ for long term storage at -75°C.

Snap-Streptavidin-WA-His (pETplasmid) was expressed in Rosettas 2 (DE3) pLysS (Merck, 71403). Culture was grown in TB medium supplemented with 30 μg/mL kanamycine and 34 μg/mL chloramphenicol, then 0.5 mM isopropyl β-D-1-thiogalactopyranoside (IPTG) was added and protein was expressed overnight at 16 °C. Pelleted cells were resuspended in Lysis buffer (20 mM Tris pH8, 500 mM NaCl, 1 mM EDTA, 15 mM Imidazole, 0,l% TritonXl00, 5% Glycerol, 1 mM DTT). Following sonication and centrifugation, the clarified extract was loaded on a Ni Sepharose high performance column (GE Healthcare Life Sciences, ref 17526802). Resin was washed with Wash buffer (20 mM Tris pH8, 500 mM NaCl, 1 mM EDTA, 30 mM Imidazole, 1 mM DTT). Protein was eluted with Elution buffer (20 mM Tris pH8, 500 mM NaCl, 1 mM EDTA, 300 mM Imidazole, 1 mM DTT). Purified protein was dialyzed overnight 4°C with storage buffer (20 mM Tris pH8, 150 mM NaCl, 1 mM EDTA, 1 mM DTT), concentrated with Amicon 3KD (Merck, ref UFC900324) to obtain concentration around 10 μM then centrifuged at 160000 ×g for 30 min. Aliquots were flash frozen in liquid nitrogen and stored at −80 °C.

### Polystyrene Beads Coating

Polystyrene beads coating with NPF was done following classical protocols (Reymann et al, MBoC 2011). 4.5 μm polystyrene beads (Polybeads Carboxylate 4.5 microns (2.6% solids-latex)) were centrifuged at 13000 g for 2 minutes on a mini spin plus Eppendorf centrifuge (Rotor F45-12-11). The pellet was then resuspended in 50 μL of a 400 nM Strep-WA solution. Beads were incubated for 15 minutes at 15°C at 950 rpm in a thermoshaker. They were then centrifuged for 2 minutes at 3800 g, resuspended in 200 uL of BSA 1% and let on ice for 5 minutes. Beads were finally centrifuged again 2 minutes at 3800 g and resuspended in 50 uL of BSA 0.1%.

### Microwells Preparation

SU8 mold with Pillars was prepared using standard protocols and silanized with Trichloro(1H,1H,2H,2H-perfluoro-octyl)silane for 1 hour and heated for 1 hour at l20°C. From the SU8 mold, a PDMS primary mold was prepared (Dow, SYLGARD 184 silicone elastomer kit) with a 1:10 w/w ratio of curing agent. PDMS was cured at 70°C for at least 2 hours. PDMS primary mold was then silanized with Trichloro(1H,1H,2H,2H-perfluoro-octyl)silane for 1 hour and heated for 2 hours at 100°C. PDMS was then poured on top of the PDMS primary mold to prepare the PDMS stamps.

Coverslips were cleaned with the following protocol: they were first wiped with ethanol (96%) and then sonicated 15 minutes in ethanol. After the first sonication, coverslips were rinced 3 times with mqH2O. They were then sonicated 30 minutes in Hellmanex 2% at 60°C.

After this second sonication, coverslips were rinsed in several volumes of mqH2O and kept in water until use. Just before use, coverslips were dried with compressed air.

For the microwells preparation, PDMS Stamps were cut in pieces and placed on the coverslips with the pillars facing the coverslip. A droplet of NOA 81 (Norland Products) was then placed on the side of the PDMS stamp and NOA was allowed to go through the PDMS Stamp by capillarity. When the NOA filled all the stamp, it was polymerized with UV light for 12 minutes (UV KUB2/ KLOE; 100% power). After polymerization of the NOA, PDMS stamp was removed and the excess of NOA was cut. Then, an additional UV exposure of 2 minutes was done and the microwells were placed on a hot plate at 110°C for at least 3 hours (or at 60°C overnight) to tightly bind the NOA to the glass.

### Lipids/SUV preparation

L-a-phosphatidylcholine (EggPC) (Avanti, 840051C) and ATTO 647N labeled DOPE (ATTO-TEC, AD 647N-161 dehydrated) were used. Lipids were mixed in glass tubes as follows: 99.75% EggPC (10 mg/mL) and 0.25% DOPE-ATTO390 (1 mg/mL). The mixture was dried with nitrogen gas. The dried lipids were incubated in a vacuum overnight. After that, the lipids were hydrated in the SUV buffer (10 mM Tris (pH 7.4), 150 mM NaCl, 2 mM CaCl_2_). The mixture was sonicated on ice for 10 minutes. The mixture was then centrifuged for 10 min at 20,238 x g to remove large structures. The supernatants were collected and stored at 4°C.

### SilanePEG30k Slides

SilanePEG (30kDa, PSB-2014, Creative PEG works) was prepared at a final concentration of 1 mg/mL in 96% ethanol and 0.1%(v/v) HCl. Slides were cleaned with the following protocol: they were sonicated for 30 minutes at 60°C in Hellmanex 2%. They were then rinced with several volumes of mqH2O. Just before use, they were dried with compressed air. Slides were plasma cleaned for 5 minutes at 80% power and directly immersed in the silanePEG solution. They were kept in the silanePEG solution until use.

### Bead Motility in microwells Assay

A typical experiment of bead motility in microwells was performed as follows. The coverslip with microwells was activated with plasma for 2 minutes at 80% power. Just after the plasma, the flow chamber was mounted with the microwells coverslip, a slide passivated with SilanePEG 30k and 180 μm height double-side tape. Lipids were then inserted in the flow chamber and incubate for 10 minutes. Lipids were then rinced with 600 μL of SUV buffer and 200 μL of HKEM buffer (50 mM KCl, 15 mM HEPES pH=7.5, 5 mM MgCl_2_, 1 mM EGTA). The reaction mix with the different proteins was then prepared and flowed in the flow cell.

A typical reaction mix was prepared with beads coated with WA (activator of the Arp2/3 complex) and 3 μM of actin monomers, 6 μM profilin, 90 nM Arp2/3 complex, 15 nM Capping Protein in HKEM Buffer and was supplemented with 0.7% BSA, 0.2% methylcellulose, 2.7 mM ATP, 5 mM DTT, 0.2 mM DABCO (motility buffer). When needed the polymerization mix was also supplement with yeast cofilin and/or cyclase associated protein (CAP).

The microwells were then closed with mineral oil (Paragon scientific Viscosity Reference Standard RTM13). The whole flow cell was then closed with VALAP and imaged under the microscope.

### Bead Motility in bulk environment

Bulk experiments were performed in a flow chamber made in the following way. A coverslip passivated with lipids or SilanePEG 30k was mounted with a slide passivated with SilanePEG 30k with double table of 70 μm height. The mix was injected in the flow chamber which was then sealed with VALAP.

### Pelleting Assay

Prior to the experiment ultracentrifuge 1.5 mL tubes were silanized with Trichloro(1H,1H,2H,2H-perfluoro-octyl)silane for 1 hour under vacuum. They were then rinsed with mqH_2_O before use.

Each individual protein of the biochemical assay (except the actin) was diluted in motility buffer in a silanized tube. At t=0 and t=24 hours, the proteins were centrifuged for 15 minutes at 55 000 rpm (rotor TLA-110). After centrifugation, supernatants were collected and pellets were resuspended in the same volume of G-buffer as the supernatant.

Samples were mixed with 4X Laemli and boiled for 5 minutes at 100°C. They were then loaded on a Stain Free Gel Mini Protean TGX 4-20% (15 wells, Biorad). Migration was performed in TGS 1X at 200V for 40 minutes. Gels were then stained with Coomassie blue. All the protein quantities were adapted to have between 0.3 and 3 μg of protein in each well of the gel.

### Imaging

Most of the experiments were done with an epifluorescence system (Ti2 Nikon inverted microscope equipped with a Hamamatsu Orca Flash 4.0 LT Plus Camera. The following objectives were used: Plan Fluor 10X DIC and S Plan Fluor ELWD 20X DIC. Time lapse were acquired with the NIS elements software (version 4.60).

Z-stacks of microwells were performed with a confocal spinning-disc system (EclipseTi-E Nikon inverted microscope equipped with a CSUXl-Al Yokogawa confocal head, an Evolve EMCCD camera (Photometrics), Plan Fluor 60X objective. Z-stacks were acquired with Metamorph software (Universal Imaging).

FRAP, k+ estimation and visualization of branched actin network formation was done on a total internal reflection fluorescence (TIRF) microscopy instrument composed of a Nikon Eclipse Ti, an azimuthal iLas^2^ TIRF illuminator (Roper Scientific), a ×60 NA1.49 TIRF objective lens and an Evolve EMCCD camera (Photometrics). Time lapse and FRAP were done with Metamorph software (Universal Imaging).

### Image analysis

Images were analyzed with FiJi (Schindelin *et al*, 2012). Data were processed with R software and plotted with GraphPad Prism. Mean and standard deviation are represented for all the data. The dot plots show the individual values with the mean and standard deviation superimposed.

Actin comets were detected with the following procedure. First, threshold was adjusted manually and images were binarized. Actin comets were detected with the Analyze particles function. Comet length was obtained with the “skeletonize” and “analyze skeleton” functions. Comet growth velocity was obtained by calculating the length difference at each time point. Tracking of comets was then done with the TrackMate plugin using the thresholding detector ((Ershov *et al*, 2022)). Comet velocity was calculated from the (x,y) coordinates obtained with the trackmate tracking. Fluorescence profiles were manually drawn on the comet tail. Bead was detected from binarization and analyze particles of the bright field movies.

#### Estimation of the ratio of actin consumed from bulk in the comet tail

To estimate the total quantity of actin in the microwell, we estimated the total fluorescence intensity. This value was constant during the time course of an experiment showing that the actin in the microwell is constant during an experiment. The quantity of actin inside a comet tail was estimated after thresholding and binarization of the comet. Those two fluorescence intensities were corrected for the fluorescence background. Then, we estimated the ratio of actin consumed from the bulk in the comet by computing the comet fluorescence over the total fluorescence of the microwell.

Half-life of bead motility was estimated by doing an exponential fit on the bead velocity curve for each independent dataset.

#### Estimation of k+

To estimate k+, we manually measured the length of actin filaments as a function of time.

### Quantitative estimations

#### Estimates of the numbers of actin molecules

The microwell is a cylinder with radius *R* = 50*μm* and height *H* = 20*μm*, so the volume of the chamber is *W* = *π R*^2^*H* ≈ 3 × (50*μm*)^2^ × 20*μm* = 1.5×10^5^ *μm*^3^. There are about 600 molecules in one cubic micron of a solution with ∼1 μM concentration (to reflect that, we use parameter *ω* ≈ 600 / (*μM* · *μm*^3^)), so the microwell contains *ω* ×*W* × 3*μm* ≈ 3×10^8^ actin subunits. About 50% of actin subunits are assembled into the actin tail in the “assembly” case, so ∼ 1.5 ×10^8^ actin subunits are in the longest tail. Thus, the total length of all filaments in the tail is 1.5 ×10^8^ × 2.7*nm* ≈ 4 ×10^5^ *μm*. Maximal actin tail length in the “assembly” case is *l* ∼ 60*μm* (Supplemental Figure 2A), so *N*_*fil*_ ∼ 4×10^5^ / 60 ∼ 6700 filaments are at every cross section of the tail. Considering that the beads are 4 microns in diameter, the cylindrical tail’s crossection area is ∼ *π R*^2^ ∼ 12*μm*^2^, the mesh size of the actin network (average distance between neighboring filaments),*ζ*, is on the order of 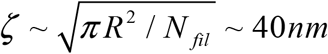, which is of the range widely reported in the literature (Kawska *et al*, 2012; Pujol *et al*, 2012).

#### Estimates of the kinetics

##### Speed

Growth speed of the actin network is *V* = *k*_*on*_*δ G*Φ, where *k*_*on*_ ≈ 10 / *μM* · *s* (Supplemental Figure 1D) is the polymerization rate, *δ* ≈ 0.003*μm* is the half-size of actin monomer, *G* is the G-actin concentration, and Φ is the dimensionless factor that accounts for geometric (filaments are not exactly parallel to the tail’s long axis), mechanical (slower growth against a mechanical load) and diffusion-limited (depletion of the local G-actin concentration at the bead-tail interface by the “consumption” of monomers by growing barbed ends) factors. Normally, parameter Φ is in the range of 0.1 - 1. Considering that the observed initial growth rate of the actin tail is ∼1 *μ*m/ min ∼ 0.02 *μ*m/s, and that 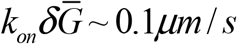 at total 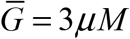, reasonable value of factor Φ ∼ 0.2 explains the data.

##### Observed kinetics of the tail growth and fraction of actin polymerized in the “assembly case

It is unlikely that the tail growth stops when critical actin concentration is reached: at the observed 50% of the assembled actin, the remaining actin concentration is too high to be critical. We therefore hypothesize that some of F-actin is not part of the actin tail but rather short actin filaments or oligomers that either polymerize near the bead and do not connect to the tail’s network, or spontaneously polymerize and then diffuse in the solute of the microwell, or both. Let *l* be the tail length, and 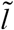 be the total length of diffuse non-tail filaments arranged into a “virtual tail” of the same geometry as the real one. The rate of filaments” assembly is *V* = *k*_*on*_*δ G*Φ. Thus, without ADF/cofilin: 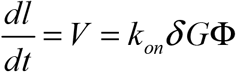. Monomer concentration is 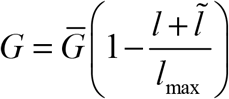 where *l*_max_ is the maximal length of the actin tail that would be achieved if *all* actin is assembled into one tail, and 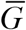 is the initial monomer concentration. Therefore,

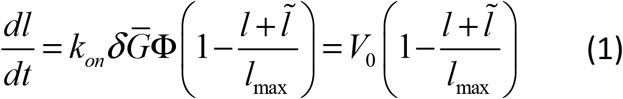

where 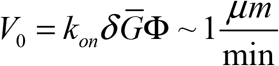 or 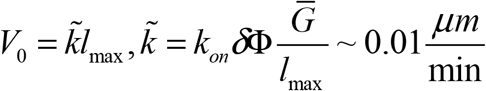

Similar equation for the non-tail F-actin is:

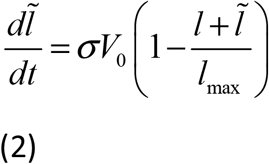

where *σ* is the parameter that accounts for the fraction of the non-tail assembly.

The solution of equation system (1-2) is: 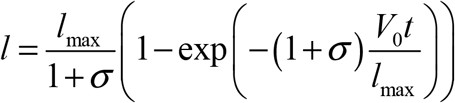and 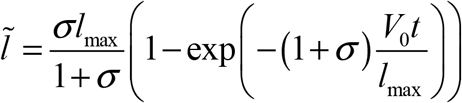

A few conclusions can be reached from these calculations:

1. the model predicts that, in the “assembly” case, the tail length grows linearly at first, and then exponentially saturates to the maximal length, as observed.
2. in this case, the order of magnitude of the time scale on which the growth speed decreases, and the tail length stabilizes is the ratio of the max tail length to the initial velocity, *T* ∼ *l*_max_ / 2*V*_0_ ∼ 100*μm* / 2*μm* / min ∼ 1 *hr*, as observed.

##### General analysis of the dynamic steady state for assembling and disassembling actin tail in the case of no aging

In this analysis we temporarily ignore the aging effect. As for the “unproductive” actin assembly, in the presence of ADF/cofilin, the short actin filaments that are not part of the tail are likely disassembled rapidly, and for simplicity, we omit the small fraction of diffuse short filaments from the analysis. We consider the following actin cycle: there is the actin tail of length *l* elongating with velocity *V* and disassembling into ADP-G-actin with rate *γ*. The net assembly flux is *V*, and the net disassembly flux is *γ l*. The equation for the tail length is:

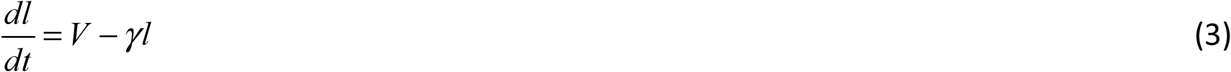

Note that area and length of the tail are proportional to each other, as the cross section area of the tail can be considered roughly constant. For convenience, in this analysis we use the tail length. ADP-G-actin concentration generated by the tail’s F-actin disassembly is *G*_*D*_ ; the disassembly flux replenishing this concentration is *flux*_*disassembly*_ = *γ l*. ADP-G-actin is recycled into ATP-G-actin, which concentration is *G*_*T*_. Respective recycling flux is *flux*_*recycling*_ = *k*_*DT*_ ×*G*_*D*_.

The dynamic equation for the ADP-G-actin concentration has the form:

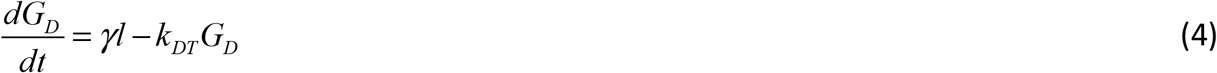

The dynamic equation for the ATP-G-actin concentration is given by the balance of the incoming recycling flux *flux*_*recycling*_ = *k*_*DT*_ ×*G*_*D*_ and outcoming assembly flux *flux*_*assembly*_ =*V* :

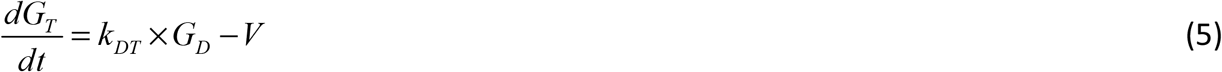

Note that according to equations 3-5 the total actin amount in the chamber, 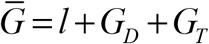, is conserved (this becomes apparent if one adds eqs 3, 4 and 5).

We measure both F-actin and G-actin concentrations in units of length. This is easy to envision in the case of the F-actin in the tail. In the case of the G-actin concentrations, there is a simple argument: if all available actin, 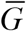, is assembled into a characteristic tail, then this tail’s length will be equal to *l*_max_. Then, any concentration *G* measured in molar can be converted into length *l* measured in microns as follows: 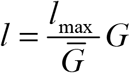. Let us now discuss the assembly rate*V*. Recall that the maximal assembly rate 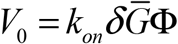, or 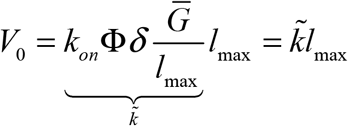. Here 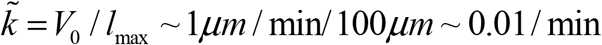. Then,

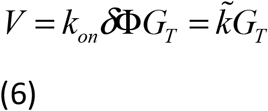

where *G*_*T*_ is now measured in units of length.

Equations 3-6 define the actin dynamics characterized by a single stable dynamic steady state, in which three *fluxes* (number of actin subunits changing chemical state per unit time) are equal and balance each other: *flux*_*disassembly*_ = *flux*_*assembly*_ = *flux*_*recycling*_. This leads to the formulas:

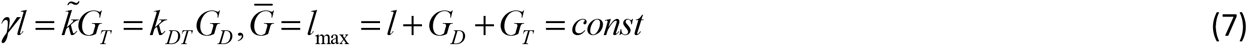

From equation 7, 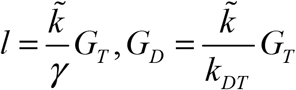. Substituting these into the conservation equation *l*_max_ = *l* + *G*_*D*_ + *G*_*T*_ = *const*, we get: 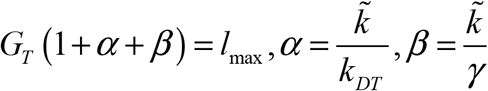. From here, we obtain:

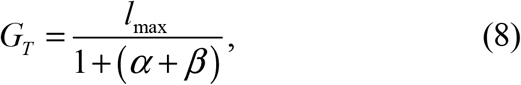

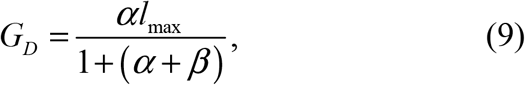

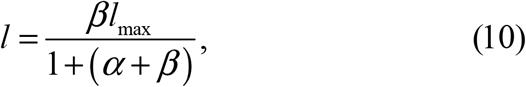

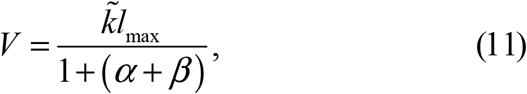

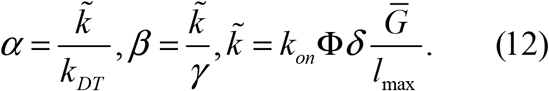

Equations (8-12) allow comparison with the data.

### Data on velocities and tail lengths in the “disassembly and “recycling cases allow rough estimates of the actin kinetics rates

In the “disassembly” case, the tail length rapidly increases at first, likely because it takes time for the disassembly machinery to start working and establishing the quasi-steady state, which happens after about two hours (Figure 2). After that, the tail length and velocity slowly (on ∼ 10 hours scale) decrease due to the aging. Thus, we use the values of the velocity, ∼ 0.5*μm* / min (Figure 2), and of the tail length, ∼ 20*μm* established a few hours after the F-actin growth starts (Supplemental Figure 2). This measured velocity is about twice smaller than the maximal velocity 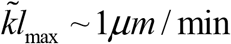 min, and according to equation 11, 1+ (*α* + *β*) ≈ 2. The measured tail length is about three-fold smaller than the maximal tail length in the assembly case, *l*_max_ ∼ 60*μm* (Supplemental Figure 2), and according to equation 10, *β* / (1+(*α*+*β*)) ≈1/ 3. From this, we deduce that *α* ∼1 / 3, *β* ∼ 2 / 3. Thus, 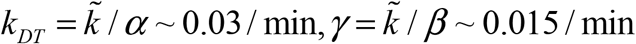 min. This means that in the “disassembly” steady state, one-third of actin is in the tail, half of it is in the ATP-G-actin form, and one-sixth of it is in the ADP-G-actin form. The tail’s F-actin is disassembled in about 1 hour, and G-actin is recycled in about 30 min (more precise estimate, see below, suggests 1 hour). This is in a good agreement with the fact that the actin tail length increases on an hour scale, and then relaxes to the quasi-steady value.

In the “recycling” case, the velocity during the first few hours is close to maximal, - 1*μm* / min (Figure 3), and the tail length is ∼ 10*μm* (Supplemental Figure 2), two hours after the start. According to equation 11, this means that1+(*α* + *β*) ≈1, and so values of both*α* and *β* are small. The measured tail length is about six-fold smaller than the maximal tail length in the assembly case, *l*_max_ ∼ 60*μm* (Supplemental Figure 3), and according to equation 10, *β* / (1+(*α* + *β*)) ≈1/ 6. From this, we deduce that *β* ∼1 / 6. Thus, 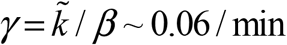 min, and so the disassembly rate increases about four-fold in the “recycling” case: the tail’s filaments disassemble, on average, in 15 min (more precise estimates that take aging effect into account suggest 10 min, see below). This agrees with the observation that the tail length in this case does not peak sharply in the beginning. The recycling rate is too fast to be estimated accurately, but it is likely to be at least an order of magnitude faster than that in the “disassembly” case, so G-actin is recycled in minutes (more precise estimates that take aging effect into account suggest that G-actin is recycled in five minutes or faster, see below). This means that in the “recycling” steady state, a small fraction of actin is in the tail, vast fraction of it is in the ATP-G-actin form, and but a tiny fraction of it is in the ADP-G-actin form.

### Fitting the data on velocities and tail lengths in the “disassembly and “recycling cases and including the aging effect into the model allows more precise estimates of the actin kinetics rates

We included the aging effect into the model by adding the aging term with respective rate*ψ* to equations 3-5, which now read:

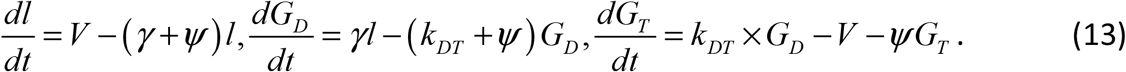

(Note that strictly speaking the tail has still to disassemble with rate *γ*, after which some monomers are aged, and some are not, but because the rate of aging is an order of magnitude slower than that of disassembly, this does not lead to a significant error). We solved the system of equations 13 numerically, and found the model parameters” values for which the fits to the data looked excellent. (The fitting procedure was ad hoc, by trying out a few tens of parameter sets.) The model results shown in the figures are the results of the numerical solutions with these optimal parameter values.

The following estimates of the parameters were obtained. In the disassembly case, the aging rate*ψ* ≈ 0.002 / min. In the recycling case, the aging rate*ψ* ≈ 0.001 / min. The F-actin disassembly rate *γ* ≈ 0.02 / min in the disassembly case, so it takes about an hour to disassemble the tail. In the recycling case,*γ* ≈ 0.12 / min, so it takes about 10 min to disassemble the tail. The recycling rate *k*_*DT*_ ≈ 0.02 / min in the disassembly case, so it takes about an hour to recycle a monomer after the disassembly. In the recycling case, any rate equal to *k*_*DT*_ ≈ 0.2 / min or faster gives a good fit, so it takes less than several minutes (could be a minute, could be seconds) to recycle a monomer after the disassembly.

## Authors contributions

**Alexandra Colin**: Conceptualization; data curation; formal analysis; investigation; visualization; methodology; writing - original draft; writing - review and editing.

**Tommi Kotila**: Resources. **Christophe Guerin**: Data curation. **Magali Orhant-Prioux**: Data curation. **Benoit Vianay**: Data curation; project administration. **Alex Mogilner**: Formal analysis. **Pekka Lappalainen**: Resources. **Manuel Thery**: Conceptualization; funding acquisition; investigation; methodology; project administration; writing - review and editing. **Laurent Blanchoin**: Conceptualization; funding acquisition; investigation; methodology; project administration; writing - original draft; writing - review and editing.

## Acknowledgements

We thank Henry N. Higgs and Tom Pollard for careful reading of the manuscript. This work was supported by the European Research Council (Consolidator Grant 771599 (ICEBERG) to MT and Advanced Grant 741773 (AAA) to LB). This work was also supported by the MuLife imaging facility, which is funded by GRAL, a program from the Chemistry Biology Health Graduate School of University Grenoble Alpes (ANR-17-EURE-0003). A.M. is supported by NSF grants DMS 2052515 and DMS 1953430. P.L is supported by a grant from the Academy of Finland (no. 302161).

## Conflict of Interest

The authors declare that they have no conflict of interest.

## Notes

### Competing Interest Statement

The authors have declared no competing interest.

### Summary of Updates

Supplemental movies added

